# GeoSinkhorn Flow: Geometry-Aware Flow Matching for Perturbation-Conditioned Dynamics in Single cell Data Phenoscapes

**DOI:** 10.64898/2026.10.16.759724

**Authors:** Ke Xu, Alistair Wilkinson, Shuang Ni, Guillaume Huguet, Mark Gerstein, Guy Wolf, Christopher J. Tape, Smita Krishnaswamy

## Abstract

Single-cell technologies increasingly enable large-scale profiling of complex biological systems across many experimental perturbations, but understanding how a system reorganizes across an entire space of conditions is difficult to interpret from single-cell embeddings alone. Phenoscapes address this by embedding entire cell-state distributions, rather than individual cells, as points in a data-driven landscape. Thus, a single dot represents one experimental condition rather than one cell, yielding a map of the perturbation or patient landscape that reveals how phenotype varies continuously across conditions. Phenoscapes remain a comparatively new and underexplored representation and existing static phenoscapes position experimental conditions relative to one another without specifying trajectories that describe how a cellular population moves from one condition to another. Flow- and trajectory-based models for single-cell data are numerous, but existing approaches operate at the level of individual cells, are not built to respect the geometry of a phenoscape, and typically cannot generalize their predicted flows to conditions not seen during training. Here we introduce GeoSinkhorn Flow, a perturbation-conditioned, geometry-aware flow-matching framework specialized designed to convert a static phenoscape into a dynamic phenoscape, in which each condition-level point is additionally connected by geodesic, condition-aware transport trajectories that are consistent with the phenoscape’s underlying geometry. GeoSinkhorn Flow can also model transitions directly between time points without constructing a phenoscape. GeoSinkhorn Flow uses heat-diffusion geometry to define manifold-aware transport costs and trains a geometric autoencoder whose latent Euclidean distances stay aligned with the static phenoscape geometry while extending it to unseen cells. By explicitly conditioning the learned transport dynamics on perturbation-specific gene or drug embeddings, GeoSinkhorn Flow generalizes its trajectories to new measurement conditions, produces intermediate progression states along a trajectory, and extends predictions to unseen cells and held-out perturbations. An unbalanced optimal transport variant further learns growth or transition weights, so that changes in relative cell-state abundance or transition propensity are modeled together with state movement. Across synthetic manifolds, patient-derived organoid drug responses, Perturb-seq knockdowns, and time-resolved CyTOF signaling, GeoSinkhorn Flow recovered intrinsic manifold geometry, inferred biologically coherent response trajectories, and generalized to held-out drug combinations, perturbation genes, and time points. Together, these results establish GeoSinkhorn Flow as a perturbation-conditioned, phenoscape-consistent dynamical framework, distinct from standard flow-matching approaches, for converting static phenoscapes into dynamic ones that generalize across experimental conditions.

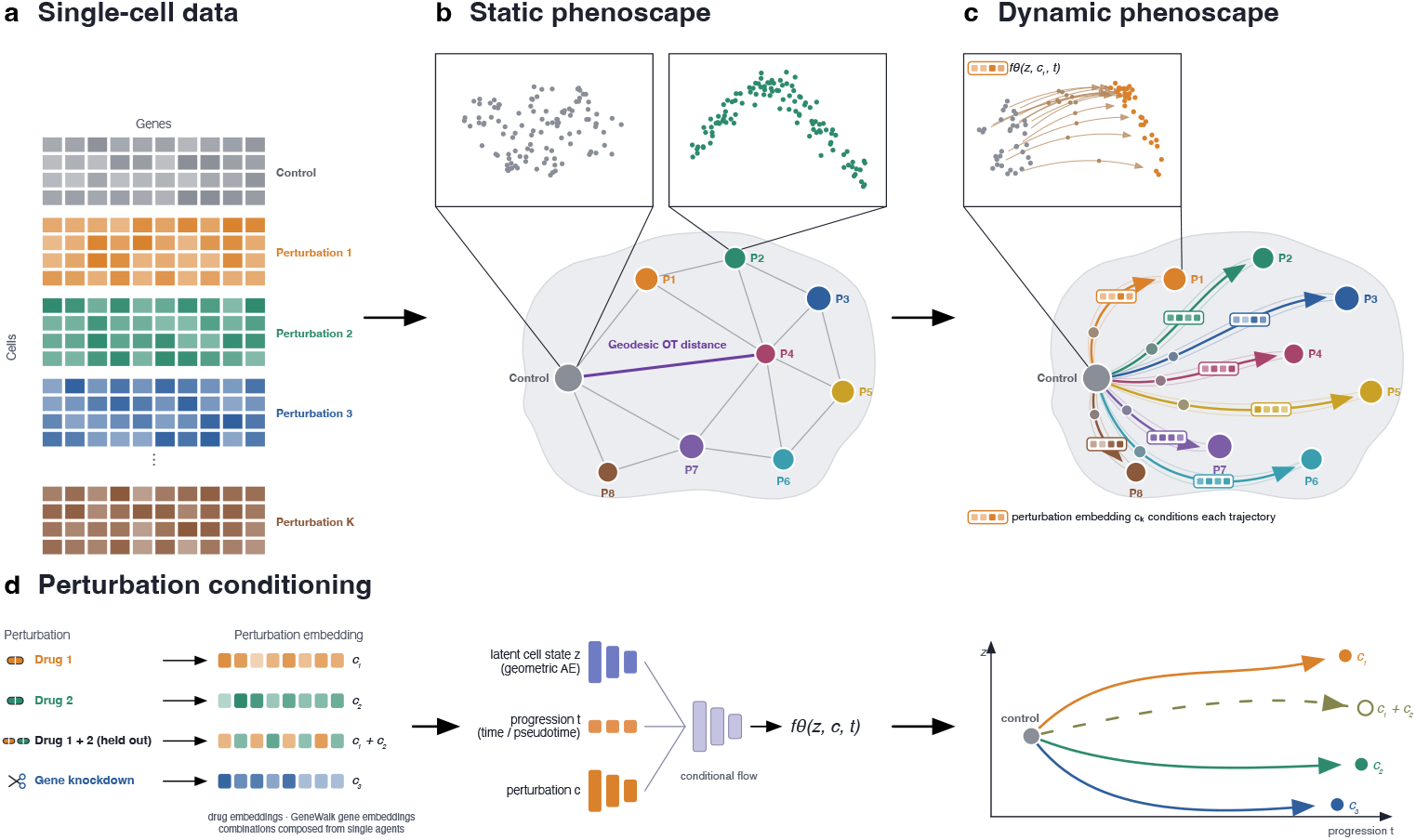

Overview of the GeoSinkhorn Flow pipeline. a) Single-cell measurements under control and perturbation conditions. b) Static phenoscape: each condition is one point representing its full cell distribution, and distances are Geodesic Sinkhorn (geodesic optimal transport) distances. c) Dynamic phenoscape: perturbation-conditioned trajectories carry the control population to each perturbation, and each trajectory is conditioned on its perturbation embedding *c*_*k*_. The inset shows single-cell trajectories under *f*_*θ*_ (*z, c*_1_, *t*). d) Perturbation conditioning: drug and gene embeddings, with combinations composed from single agents, condition a flow over latent cell state *z* and progression *t*. This lets a single model predict a distinct trajectory for each perturbation, including held-out combinations (dashed).

## 1 Introduction

A fundamental challenge in biology is understanding how a complex biological system reorganizes across an entire space of perturbations, such as drug treatments, genetic knockdowns, patient states, not just how one cell responds in isolation. Single-cell technologies now profile millions of cells across thousands of such conditions, but a collection of single-cell embeddings does not by itself show how the system as a whole moves through the space of conditions. Neither does it show which conditions are phenotypically close to one another.

A phenoscape addresses this directly: it embeds entire cell-state distributions, rather than individual cells, as points in a shared landscape. The distances between points correspond to a transport distance between the underlying populations. One dot represents one drug, genetic perturbation, or patient sample summarized from its cell population. [1] provide an early example, embedding colonic organoid cultures by their Earth mover’s distance profiles to map how oncogenic and microenvironmental cues shape stem cell fate A phenoscape can be constructed from the geometry-aware transport distances between cell populations calculated by Geodesic Sinkhorn [2]. The result is a map of the perturbation or patient landscape, showing how phenotype varies continuously across experimental conditions in a way no single-cell embedding can.

Phenoscapes are a recent and still underexplored representation of single-cell perturbation data. Existing work has focused primarily on their static organization: static phenoscapes reveal which perturbations produce similar cellular distributions, but not how populations transition between conditions or through which intermediate states. Exploring these dynamics would extend the phenoscape from a descriptive map into a model of biological response, capable of representing continuous trajectories illustrating changes in population composition, and responses to previously unseen conditions.

Constructing a phenoscape requires reducing each unpaired, high-dimensional cell-state distribution to a point while preserving the geometry of the space over which its cells lie. Optimal transport (OT) provides a natural framework for comparing such distributions. Although OT does not inherently require Euclidean geometry, its ground cost is commonly defined using Euclidean distances, which can be poorly suited to single-cell data supported on curved, low-dimensional manifolds: straight-line distances may create shortcuts through sparsely sampled or biologically unsupported regions.

Diffusion geometry provides an alternative by using heat propagation on a neighborhood graph to approximate intrinsic manifold distances. Methods such as Diffusion Maps, PHATE, and HeatGeo use this principle to represent the geometry of individual cells; in particular, HeatGeo constructs an embedding that preserves heat-geodesic dissimilarities derived from the graph heat kernel [3]. Geodesic Sinkhorn lifts the same geometric principle from individual cells to distributions by using heat-kernel-derived ground geometry in entropy-regularized OT [2], yielding geometry-aware distances and couplings between cell populations. HeatGeo and Geodesic Sinkhorn thus play complementary roles: the former represents heat-diffusion geometry at the cell level, whereas the latter uses that geometry to compare population-level distributions. However, these graph-derived distances and couplings remain tied to the observed cell populations: they do not define continuous trajectories or extend naturally to unseen cells. Modeling phenoscape dynamics therefore requires transferring this diffusion geometry into a continuous representation in which population flows can be learned.

Developing such a dynamic representation requires a flow model that preserves the geometry used to construct the phenoscape. We introduce GeoSinkhorn Flow, a flow-matching framework that extends a static Geodesic-Sinkhorn phenoscape into a continuous and predictive dynamical system. GeoSinkhorn Flow trains a geometric autoencoder whose latent Euclidean distances remain consistent with the heat-diffusion geometry underlying the phenoscape, allowing trajectories learned in this space to follow the same manifold structure used to organize the static map. Geodesic Sinkhorn couplings provide geometry-aware supervision for local population transitions, which conditional flow matching integrates into continuous, condition-aware trajectories. This construction enables generalization to unseen drug combinations, perturbation genes, and intermediate time points. An unbalanced OT extension further models changes in population mass alongside cell-state movement.

We first validated GeoSinkhorn Flow on synthetic manifolds, where its learned latent distances accurately recovered ground-truth geodesic geometry. We then applied it to three biological systems. In patient-derived colorectal cancer organoids, GeoSinkhorn Flow reconstructed patient-specific drug-response trajectories and generalized from single agents to unseen combinations. In a CAD Perturb-seq screen, it organized knockdowns by their downstream transcriptional responses, reconstructed continuous control-to-perturbation dynamics, and generalized to unseen held-out gene. In time-resolved CyTOF data, it recovered continuous, treatment- and genotype-dependent signaling trajectories from discrete measurements. Together, these results establish GeoSinkhorn Flow as a framework for transforming cross-sectional and longitudinal single-cell data into predictive, geometry-aware dynamic phenoscapes.

## 2 Results

### 2.1 GeoSinkhorn Flow Method overview

GeoSinkhorn Flow is designed to transform a static phenoscape constructed from unpaired single-cell measurements under control and perturbed conditions into a continuous, geometry-aware dynamic phenoscape. Perturbations, such as drug treatments or genetic knockdowns, are represented by perturbation embeddings that condition the learned population dynamics. The goal is to connect condition-level population states through continuous trajectories, while using cell-level transport paths to describe how each population reorganizes between conditions. These trajectories preserve the intrinsic geometry of the data and generalize to new cells or perturbations. Figure 1 summarizes the full GeoSinkhorn Flow pipeline, and full methodological details are provided in the Method Section 4.

**Fig. 1.**
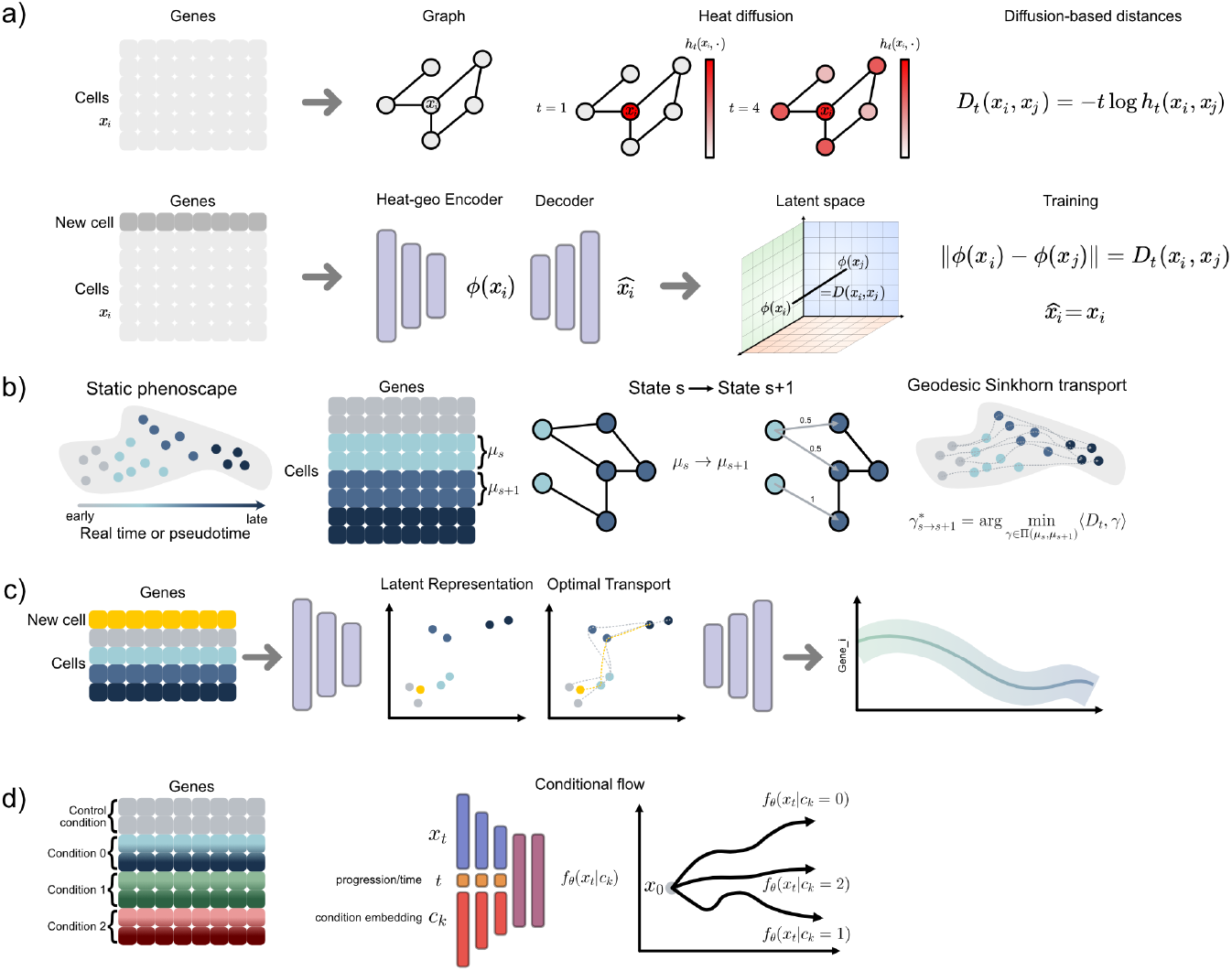
Overview of the GeoSinkhorn Flow pipeline. a) we construct a weighted nearest-neighbor graph and use heat diffusion on this graph to define a diffusion-based distance *D*_*t*_ between cells. Because this graph-based distance does not directly generalize to new observations, we train an autoencoder so that Euclidean distance in the latent space matches *D*_*t*_ while still allowing reconstruction in the original feature space. b) we use real time or pseudotime to define a progression coordinate on the static phenoscape, and compare neighboring population states along this progression. These states are encoded into the geometry-aligned latent space, where OT is computed and decoded back to the ambient space to define Geodesic Sinkhorn transport between successive states. By combining these local transitions, we obtain a geometry-aware dynamic flow on the phenoscape. c) new cells are encoded into the geometry-aligned latent space, where Euclidean OT provides a tractable approximation of geodesic transport because the latent representation has been trained to preserve diffusion distances. The decoder then maps transported latent states back to the ambient space, enabling interpretation of treatment effects and population transitions in the original cellular coordinates. d) we train a conditional flow over the progression variable using neighboring population states and treatment embeddings. This allows the same initial cell state to evolve toward different endpoints under different conditions and at different progression times, turning the static phenoscape into a navigable dynamical system.

GeoSinkhorn Flow proceeds in three stages:

1. It estimates the geometry underlying the observed phenoscape by constructing a weighted nearest-neighbor graph and using heat diffusion on that graph to define a diffusion-based distance between cells.
2. A geometric autoencoder is trained so that Euclidean distances in latent space approximate these diffusion distances while still allowing reconstruction in the original feature space because graph distances do not naturally extend to unseen observations. This learned latent space provides a geometry-aligned representation in which optimal transport between source and target populations can be computed efficiently.
3. The transport relations between neighboring progression states are elaborated into a conditional flow model that takes as input a cell state, a progression variable, and a treatment embedding, and outputs a continuous cell-state trajectory through latent space.

Here, the progression variable represents the position of a cell along the biological response, and may correspond to measured experimental time when real time points are available, or to an inferred ordering such as pseudotime or MELD-based control-to-treatment progression when the data consist only of cross-sectional snapshots. Collectively, these cell-level trajectories describe the continuous evolution of a population between condition-level points in the phenoscape. In this way, the model converts static population relationships into a perturbation-conditioned flow that can be evaluated at arbitrary intermediate progression values and transferred to unseen perturbations. At inference, new cells are encoded into the latent space, evolved according to the learned dynamics for a given perturbation and progression value, and then decoded back into the original feature space.

This structure is important for the biological applications considered here. The geometric stage encourages that cell-level transitions follow the manifold traced by real cellular states rather than straight lines in ambient measurement space. The transport stage links source and target populations in a way that is consistent with this geometry, and the resulting transport relations are used to supervise the dynamic model. When used in its unbalanced form, this transport stage additionally models changes in population mass through learned growth rates. This unbalanced transport allows the GeoSinkhorn Flow framework to capture not only state transitions but also expansion or depletion of cell populations under perturbation. The final conditional flow stage then turns these geometry-aware correspondences into predictive trajectories connecting condition-level points, enabling continuous inference of treatment responses, interpolation of missing states, and extrapolation to new conditions within a shared dynamic phenoscape.

### 2.2 Validation of geometry-aware distance recovery on synthetic manifolds

We first evaluated whether the geometric autoencoder in GeoSinkhorn Flow learns a latent space whose Euclidean distances recover the geometry-aware distance structure of the original data manifold. This is a key requirement of the model: graph-based distances computed from samples in the original space capture the intrinsic geometry of the manifold, but they are tied to a fixed observed point cloud and do not directly define distances for unseen observations. The autoencoder is therefore trained to map observations into a latent representation where standard Euclidean distances approximate these graph-based geometry-aware distances, allowing downstream optimal transport and flow matching to be performed in a continuous latent space.

To test this distance recovery in a setting where the ground truth is known, we generated three synthetic manifolds: a Swiss roll, a saddle surface, and a toroidal surface. These datasets contain different types of nonlinear geometry, including curvature, folding, and periodicity. For each dataset, samples were organized into local distributions on the manifold, and ground-truth geodesic distances were computed analytically or from the underlying manifold coordinates as described in Section 4.8.1. This provides a controlled validation setting in which we can compare distribution-level distances induced by ambient Euclidean costs, graph-based geometry-aware costs, and Euclidean distances in the learned latent space.

Figure 2a-b illustrates the motivation for this experiment. In the original ambient space, Euclidean distances can create shortcuts across unsupported regions of a curved manifold, whereas geodesic distances follow the sampled surface. This difference affects not only point-to-point distances, but also Wasserstein distances between distributions supported on the manifold. Across the three synthetic manifolds, the learned latent distances show high agreement with the ground-truth geodesic Wasserstein distances and improve over the ambient Euclidean baseline (Figure 2c,d). A detailed description of the distribution-level distance comparisons, evaluated methods, and quantitative metrics is provided in Appendix B.2.2. These results support the central geometric assumption of GeoSinkhorn Flow : Euclidean distances in the learned latent space can serve as geometry-aware ground costs for downstream transport and flow learning.

**Fig. 2.**
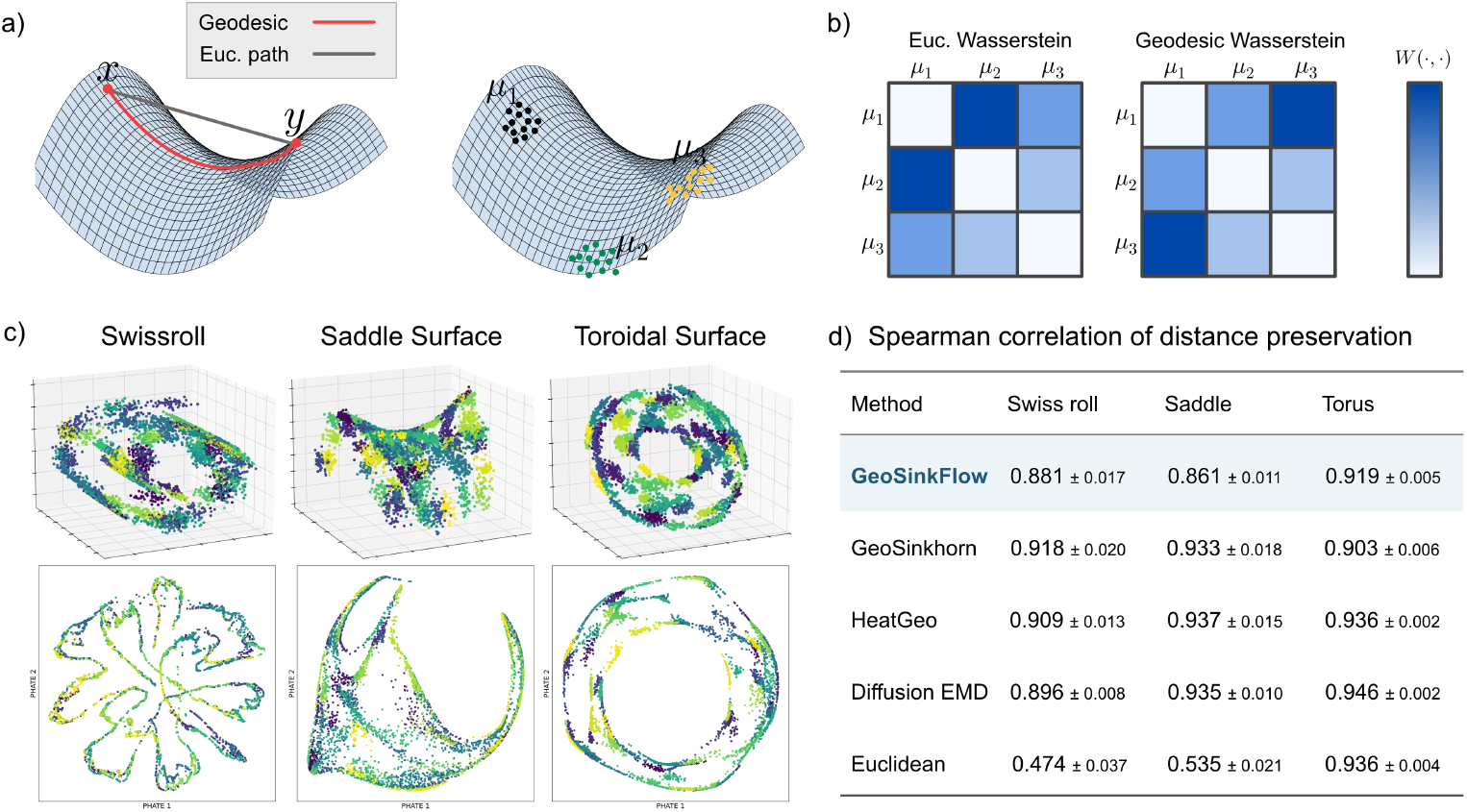
Synthetic manifold experiments validate geometry-aware distance recovery in latent space. a) Comparison of geodesic and Euclidean paths between two points on a curved manifold, illustrating how Euclidean distances can shortcut through ambient space rather than follow the data geometry. b) Example heatmaps comparing Euclidean and Geodesic Wasserstein distances between distributions. c) Synthetic Swiss roll, saddle surface, and toroidal surface datasets in 3D plots, with corresponding PHATE embeddings showing their recovered manifold structure. d) Quantitative comparison of distance methods against ground-truth geodesic distances across the three manifolds, evaluated by Spearman correlation; full metrics are reported in Supplementary Table B2. The learned model recovers the intrinsic distance structure more accurately than the ambient Euclidean baseline, supporting the use of latent Euclidean distances as geometry-aware distances for downstream transport and flow learning.

### 2.3 GeoSinkhorn Flow generalizes to unseen drug combinations in patient-derived organoids

We evaluated GeoSinkhorn Flow on a single-cell mass cytometry dataset of colorectal cancer patient-derived organoid (PDO) cultures under drug perturbation [4]. In this dataset, each sample is an unpaired snapshot of cells measured under either a control or a drug-treatment condition. The PDO setting therefore tests whether GeoSinkhorn Flow can infer a continuous treatment-response process from cross-sectional single-cell measurements. Details of data acquisition and preprocessing are provided in Section 4.8.2.

For these experiments, monoculture PDOs with drug treatments were used. We trained the model only on single-treatment conditions and withheld combination-treatment conditions as unseen test cases. This defines a stringent held-out generalization task: given control and single-condition responses, the model must infer the trajectories induced by combinations that were never observed during training. Successful generalization therefore means that the predicted combination responses form coherent trajectories in latent space and recover the major growth and marker-level trends of the held-out combinations.

In this non-longitudinal setting, we used MELD scores [5] computed for each control-to-treatment comparison to define pseudotime bins. MELD estimates a smooth likelihood of each cell belonging to the control or treatment condition over the single-cell manifold, thereby providing a continuous score that orders cells from control-like to treatment-like states (Supplementary Figure B2b). Each treatment condition was represented as an embedding, and combination treatments were constructed compositionally from the embeddings of their constituent single agents, enabling generalization from observed single-treatment responses to unseen combination perturbations; this compositional scheme is illustrated in Supplementary Figure B2a.

Phenoscapes of PDOs 21, 23 and 27 were constructed by aggregating interpolated cell trajectories produced by GeoSinkhorn Flow into intermediate distributions between the control-like and treatment-like cells. Paired Trellis was used to embed the trajectories into condition level space [4]. The controls are anchored together to the origin while condition level trajectories capture the treatment effect of the perturbation (Figure 3a). Next, we summarized treatment relationships at the population level using pairwise Spearman correlations between treatment trajectories and Ward hierarchical clustering (Figure 3b-c). This provides population-level validation that GeoSinkhorn Flow captures biologically meaningful response structure while preserving the heterogeneity of treatment effects across PDOs. Treatments show clear divergence from their respective baseline controls with patient-specific responses.

**Fig. 3.**
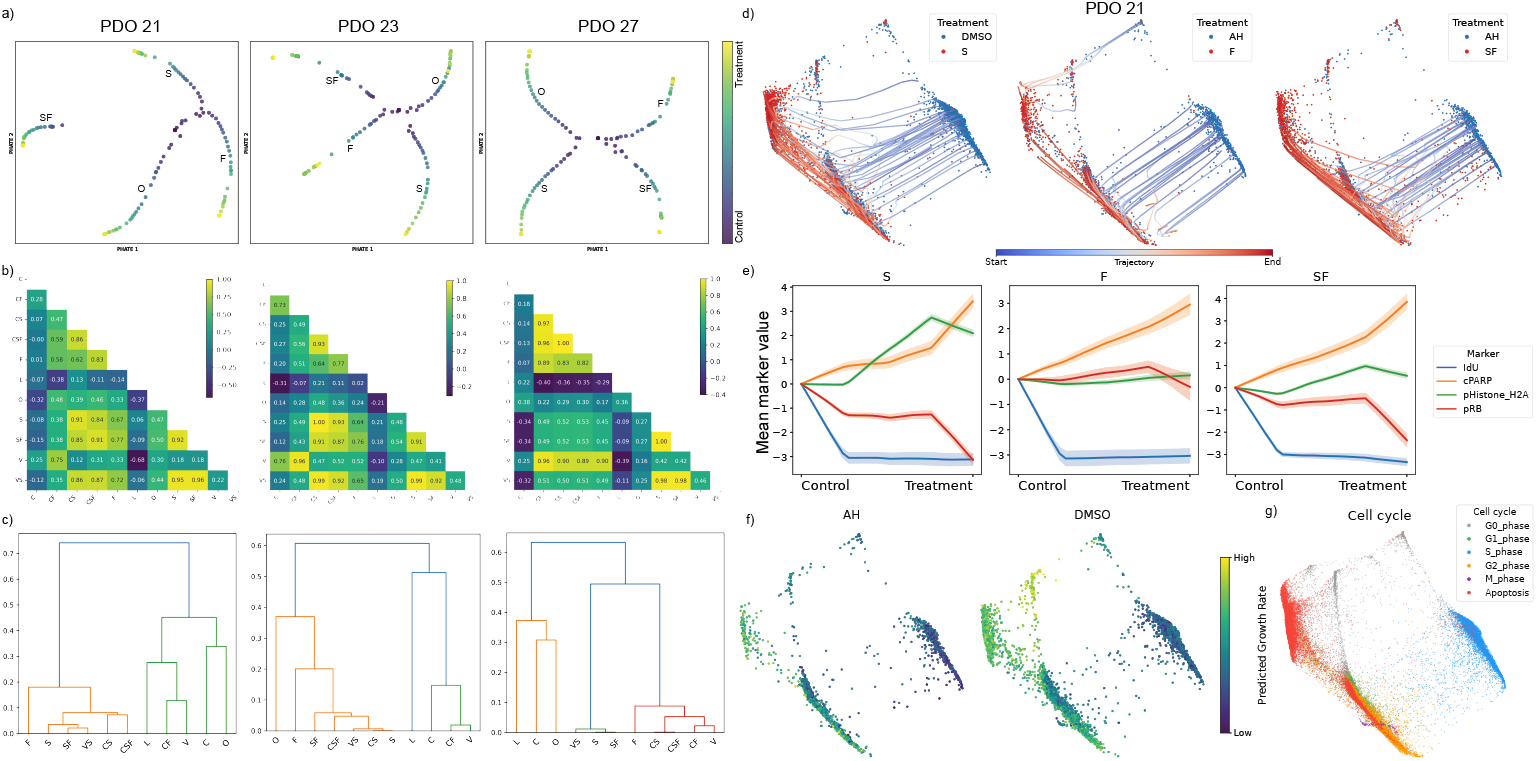
GeoSinkhorn Flow captures control-to-treatment PDO phenoscape transitions and generalizes to unseen drug combinations. a) For PDOs 21, 23 and 27, phenoscape embeddings of S, F, O and SF with condition resolution trajectories from control to perturbation b) Pairwise Spearman correlations between treatment trajectories across IdU, cPARP, pHistone-H2A and pRB. c) Dendrogram from Ward linkage on the same Spearman dissimilarity matrix. d) Inferred cellular trajectories in PHATE space for single treatments (S, F), and combination treatment (SF). e) Predicted marker-level trajectories for single treatments (S, F) and held-out combination (SF) evaluated on apoptosis and cell-cycle markers (pRB, cPARP, pHistone H2A, IdU). Values show mean marker changes relative to the control condition at *t* = 0. f) Predicted growth rate across control cells over PHATE. g) Cell-cycle phase over PHATE, assigned by marker-intensity gating as described in Supplementary Figure B1.

In Figure 3a, condition-level trajectories for S, O, F and SF diverge from the controls centered at the origin. Each trajectory captures the shift in the cellular composition induced by its corresponding treatment. The different trajectory directions and position highlights the known patient-specific treatment effect of each drug. SN-38-containing combinations (CS, CSF, SF, VS) produced aligned trajectories across all three PDOs (Figure 3b), with pairwise Spearman correlations frequently exceeding *ρ >* 0.85. SN-38 acts as the dominant driver, trapping topoisomerase I-DNA cleavage complexes to induce replication-fork collapse and S-phase double-strand breaks. Cetuximab and berzosertib contributed a negligible treatment effect while 5-FU intensified, rather than redirected, the response by stalling replication forks through thymidylate synthase inhibition. Thus, SN-38-containing combinations had similar treatment effects and high Spearman correlation. Finally, Ward hierarchical clustering reproduced a two-group structure in all three PDOs, with S-backbone treatments merging at low linkage distances (*<* 0.1) (Figure 3c).

The resulting trajectories then trace smooth paths through PHATE space for both observed single treatments (S, F) and held-out combination (SF) (Figure 3d). The trajectories flow from S phase to apoptosis, accurately capturing how SN-38, 5-FU and their combination shift S-phase cells toward an apoptotic state through a DNA-damage-associated response. At the marker level, the predicted PDO21 trajectories remain coherent with this expected treatment-response trend. SN-38 (S), Oxaliplatin (O) and 5-FU (F) capture a canonical apoptosis signature with increasing cPARP suggesting caspase-3 is active and executing apoptosis while IdU and pRB decrease, implying cells have ceased DNA synthesis due to exiting the cell cycle (Figure 3e; Supplementary Figure B2). The predicted growth-rate landscape varies across control cells, with higher growth scores concentrated in regions enriched for apoptotic cells (Figure 3f-g). Cell-cycle states were assigned by marker-intensity gating using cPARP, Cyclin B1, IdU, pHH3, pHistone H2A and pRB; details of the gating strategy and thresholds are provided in Supplementary Figure B1.

The same analysis on PDO23 and PDO27 shows that GeoSinkhorn Flow preserves coherent treatment-response trajectories while capturing stronger patient-specific heterogeneity in marker-level dynamics (Supplementary Figures B3 and B4). Pearson and Spearman correlation analysis of trajectory start and end points for seen single-drug and unseen combinatorial treatments on the PDO 21 dataset is provided in Supplementary Figure B5. Together, these results show that GeoSinkhorn Flow learns a continuous treatment-response flow that can be evaluated at intermediate states and generalized beyond the perturbations observed during training.

### 2.4 GeoSinkhorn Flow reconstructs continuous transcriptional responses and generalizes to unseen genetic perturbations

We next asked whether GeoSinkhorn Flow could reconstruct continuous transcriptional responses from static genetic perturbation data and transfer the learned dynamics to perturbations excluded from training. We applied the model to a Perturbseq dataset in which CRISPR interference was used to target coronary artery disease (CAD)-associated genes in human arterial endothelial cells [6]. The original study connected CAD genetic risk to endothelial transcriptional programs, including convergence of multiple risk-associated genes on CCM signaling. Because each knockdown was measured only at a cross-sectional endpoint, this dataset provides a stringent setting in which to test whether a dynamic response landscape can be inferred without observing intermediate states.

We first examined the population-level organization of the perturbations. Pairwise geometry-aware Sinkhorn distances between batch-corrected knockdown responses were embedded into a two-dimensional perturbation phenoscape (Figure 4a). Because these distances were calculated from the resulting cellular distributions, proximity in this phenoscape reflects similarity between downstream transcriptional responses rather than similarity between the targeted genes themselves. Negative-control and safe-targeting perturbations occupied a control-proximal region, whereas CAD-associated perturbations formed several distinct response neighborhoods. These neighborhoods included perturbations associated with endothelial junction and signaling programs, such as *CCM2, CDH5, FLT1*, and *SMAD3* ; vasoactive and angiocrine responses involving *NOS3, EDN1*, and *PDGFB* ; vascular-remodeling responses involving *KRIT1, KLF4*, and *ENG* ; and matrix- or cytoskeleton-associated responses involving *FN1, POSTN, RHOA*, and *CDC42*. Thus, the static phenoscape organized perturbations by their realized endothelial phenotypes and provided the population-level geometry that GeoSinkhorn Flow subsequently resolved into continuous response trajectories.

**Fig. 4.**
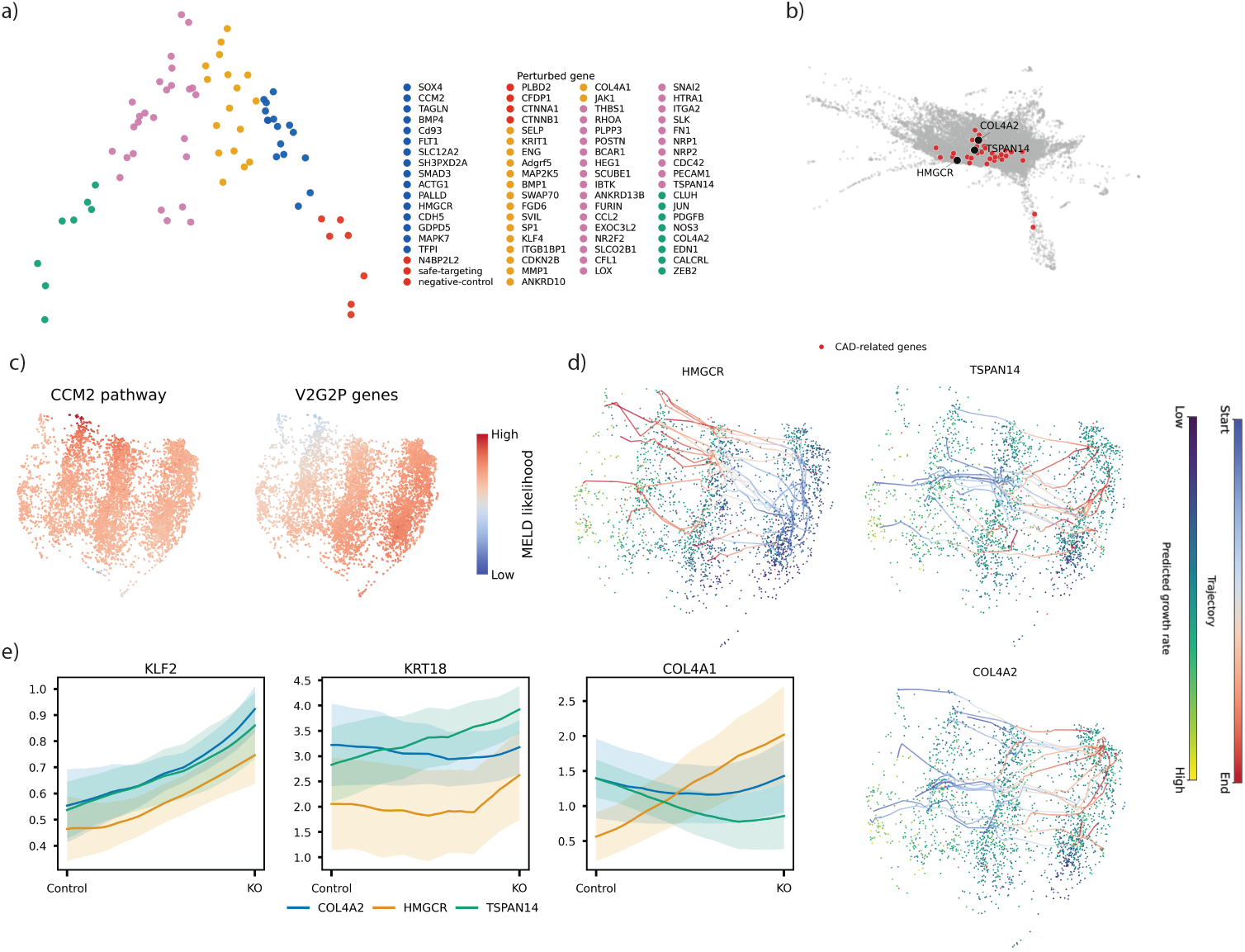
GeoSinkhorn Flow models geometry-aware transcriptional response trajectories in Perturb-seq. a) Perturbation phenoscape computed from pairwise geometry-aware Wasserstein distances between batch-corrected knockdown effects. Each point represents one perturbed gene, and colors denote clusters of perturbations with related transcriptional responses. b) PHATE visualization of the GeneWalk embedding space used to condition the model on perturbation identity. CAD-related perturbation genes are highlighted, with the representative trajectory genes labelled. c) Cell-level PHATE embeddings colored by MELD likelihood for the CCM2 pathway and V2G2P CAD gene program. These maps show where CAD-associated perturbation programs are enriched across the transcriptomic manifold. d) Predicted GeoSinkhorn Flow trajectories for representative CAD-related knockdowns projected onto the cell-level PHATE embedding. Source cells are colored by predicted growth rate, and inferred paths are colored from trajectory start to endpoint. *HMGCR* and *COL4A2* provide training-reference examples, whereas *TSPAN14* evaluates generalization to a held-out perturbation gene. e) Predicted expression trajectories for selected response transcripts across the same perturbations. Lines show mean predicted expression along the inferred control-to-knockdown trajectory, and shaded regions show variability across sampled trajectories.

To determine whether these population-level relationships were reflected within the cellular manifold, we mapped CAD-associated response programs onto the cell-level PHATE representation. MELD likelihoods associated with the CCM2 and V2G2P CAD programs varied smoothly across the manifold rather than separating cells into discrete endpoint groups (Figure 4c). This continuous organization indicates that the effects of CAD-related perturbations span graded transcriptional states. It also supports the use of MELD-derived control-to-knockdown coordinates to order cells along each perturbation response when experimentally measured intermediate states are unavailable.

Using these perturbation-specific progression coordinates, GeoSinkhorn Flow reconstructed continuous trajectories from control-like states toward knockdown-enriched regions of the cellular manifold (Figure 4d). We examined *HMGCR* and *COL4A2* as perturbations represented during training and *TSPAN14* as a held-out perturbation. These examples capture distinct aspects of CAD-associated endothelial biology, including lipid regulation, basement-membrane organization, and vascular signaling. For both training and held-out perturbations, the inferred trajectories remained within densely populated regions of the PHATE manifold and followed the corresponding control-to-knockdown MELD gradients. Pair-specific analyses confirmed that the trajectories progressed toward states enriched for the appropriate perturbation response (Supplementary Figure B6). Importantly, the coherent trajectory inferred for held-out *TSPAN14* shows that conditioning on biologically informed GeneWalk embeddings allowed the model to transfer response dynamics to a perturbation that was not used to train the flow.

The inferred trajectories also provided a continuous view of transcriptional regulation that was not available from the original endpoint comparisons. Predicted expression profiles revealed coordinated changes in endothelial flow-response and extracellular-matrix genes, including *KLF2* and *COL4A1*, together with changes in the state-associated transcript *KRT18* (Figure 4e). These expression curves linked movement through the cellular manifold to specific molecular programs and distinguished the responses induced by different knockdowns. In parallel, the unbalanced transport component assigned perturbation-dependent mass-change scores across the source population, identifying cellular regions predicted to undergo relative expansion or depletion during each response.

Together, these analyses extend the static perturbation phenoscape into a dynamic representation of CAD-associated endothelial responses. The population-level map identifies perturbations that produce similar transcriptional states, whereas the learned flows resolve how cells move toward those states and which gene programs change along the way. The successful reconstruction of the held-out *TSPAN14* response further demonstrates that GeoSinkhorn Flow can generalize continuous dynamics across genetically distinct but biologically related perturbations.

### 2.5 GeoSinkhorn Flow learns continuous signaling dynamics from time-resolved CyTOF signaling data

As a time-resolved signaling benchmark, GeoSinkhorn Flow was applied to a CyTOF dataset from HeLa spheroids carrying wild-type *PIK3CA* or one or two endogenous copies of the oncogenic *PIK3CA*^H1047R^ allele [7]. In the original study, these cells were stimulated with EGF or IGF1 to determine how oncogenic PI3K*α* reshapes growth-factor signaling specificity. The dataset provides a stringent test for dynamic modeling because signaling responses vary jointly with time, ligand identity and *PIK3CA* allele dosage. Details of data acquisition and preprocessing are provided in Section 4.8.4.

This setting tests whether discrete CyTOF snapshots can be converted into continuous signaling trajectories across genotype and stimulation conditions. We first summarized the generated multidimensional CyTOF profiles in a phenoscape, constructed by applying PHATE [8] to marker-wise EMD summaries of the predicted response distributions (Figure 5a). For each generated genotype–treatment–time condition, an Earth Mover’s Distance (EMD; Wasserstein distance) score was calculated for each of the five EMD-PHATE signaling markers relative to the corresponding single-cell marker distribution in the WT H_2_O reference condition. These marker-wise EMD scores quantify how far each generated response distribution shifts from the WT control state and define a condition-by-marker response matrix. The resulting EMD-PHATE space separates H_2_O control-like states from growth-factor-stimulated states and organizes EGF and IGF1 responses along time-ordered transitions with visible genotype-dependent structure, consistent with the condition-level signaling organization reported in the original study.

**Fig. 5.**
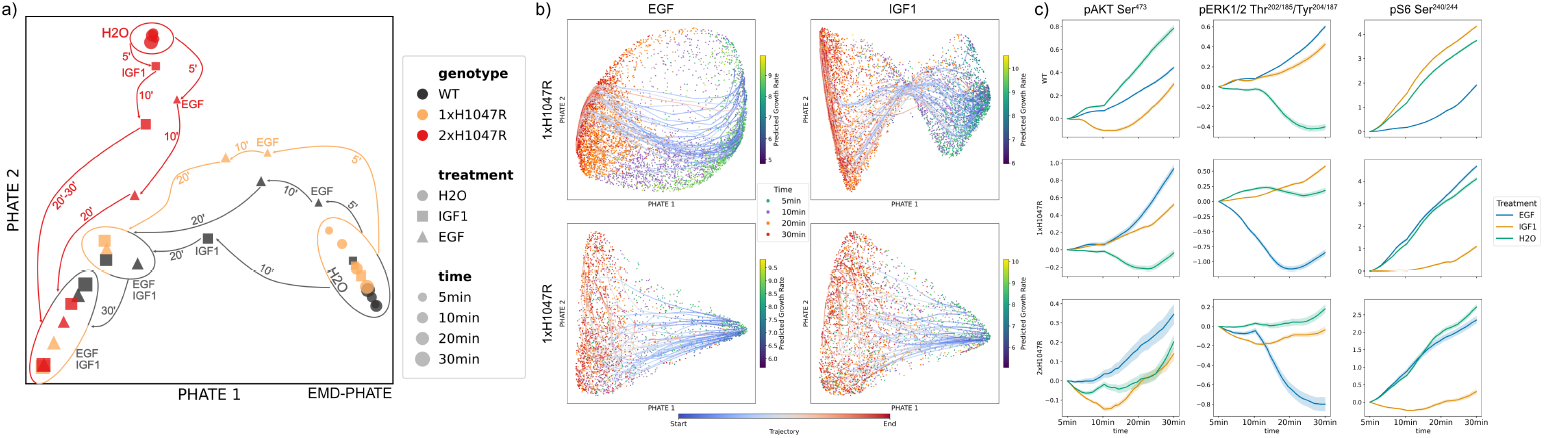
Continuous inference of signaling trajectories from discrete time-point CyTOF measurements in PIK3CA-mutant HeLa spheroids under EGF and IGF1 stimulation with H_2_O control. Mass cytometry (CyTOF) data were collected from cycling, non-apoptotic HeLa spheroid cells expressing wild-type (WT) PIK3CA, or one (1×H1047R) or two (2×H1047R) copies of the oncogenic *PIK3CA*^H1047R^. a) Generated signaling EMD-PHATE: Phenoscape of generated genotype– treatment–time distributions, computed from marker-wise EMD scores relative to the WT–H_2_O reference condition. The embedding separates control-like and growth-factor-stimulated states while preserving time-ordered, genotype-dependent response structure. b) PHATE single-cell embedding reveals continuous inferred trajectories across time in 1×H1047R and 2×H1047R cells under EGF and IGF1 stimulation. c) Marker-level trajectories show generated temporal responses for representative signaling markers across WT, 1×H1047R and 2×H1047R genotypes under EGF, IGF1 and H_2_O conditions; the 20 min measurements were withheld during training.

The generated dynamics were also examined in a cell-state embedding. In the 1×H1047R and 2×H1047R populations, predicted paths traverse the PHATE space across the measured time interval under both EGF and IGF1 stimulation (Figure 5b). The trajectories remain organized by time and condition, indicating that the model produces coherent cell-level paths through the signaling manifold rather than disconnected endpoint predictions. Collectively, these paths explain how the population evolves between the condition-level time points in the dynamic phenoscape. Because this short signaling time course captures acute pathway activation rather than changes in cell abundance, the weights shown on source cells are interpreted as transition weights: higher values indicate signaling states that more readily progress toward the next measured response state.

At the marker level, the generated trajectories show condition-specific temporal responses across PI3K/MAPK readouts, including pAKT, pERK1/2 and pS6 (Figure 5c). The distinct trajectory shapes across genotype–ligand conditions show that the model retains allele-dose- and stimulus-dependent signaling patterns rather than averaging them into a common response. These curves summarize how the model represents each genotype–ligand condition as a continuous response profile rather than as independent time-point measurements, indicating that GeoSinkhorn Flow preserves coordinated temporal signaling relationships in the generated populations. To assess temporal generalization, the 20 min measurements were withheld during training and predicted from the remaining time points. Agreement at the held-out time point was evaluated by comparing generated and original single-cell marker distributions using marker-wise Wasserstein distances. Additional marker trajectories are provided in Supplementary Figure B7a, and the distribution-level evaluation is reported in Supplementary Figure B7b.

## 3 Discussion

In this work, we introduced a generalizable method that uses diffusion-based distances and conditional flow matching in a learnt latent space to construct continuous dynamic phenoscapes that capture heterogeneous phenotypic states and model trajectories in complex biological data. A major contribution of our method is moving from a diffusion-based static OT to a learnable latent space via a geometric autoencoder, which generalizes the diffusion-based distance structure to previously unseen observations and supports OT-coupled conditional flow matching in a geometry-aligned latent space. Existing methods often either preserve geometry without modeling continuous dynamics, or learn continuous dynamics without explicitly preserving manifold geometry. Across single-cell mass cytometry, single-cell transcriptomic perturbation and time-resolved single-cell signaling datasets, this framework facilitated accurate interpolation of intermediate time points and prediction of novel treatment responses using continuous flows between distributions, thus producing dynamic trajectories between phenotypic states in a phenoscape embedding.

We applied GeoSinkhorn Flow across datasets with distinct modalities and experimental structures, including patient-derived organoid (PDO) drug perturbation data, Perturb-seq transcriptomic perturbation data and time-resolved CyTOF signaling data. In each setting, the method was able to use either measured time or inferred progression to learn continuous transitions between neighboring states while preserving the underlying geometry of the data. The PDO data showcase the potential and complexity of modeling individualized treatment dynamics. Each PDO exhibits changes in marker expression under drug perturbation that are successfully captured in GeoSinkhorn Flow ‘s patient-specific dynamics while sharing a common learned latent space. This shared manifold with patient-specific dynamics mirrors the tumor heterogeneity within CRC, providing a concrete approach to pool geometric information across patient data without conflating their patient-specific drug responses. Furthermore, the model predicts trajectories of unseen drug combinations from single-drug training data, with trajectories spanning phenotypes represented in both control and treatment conditions. This is crucial as combination therapies represent the standard of clinical care and deriving all pairwise or higher order conditions experimentally would be time-consuming and expensive. In this setting, geometry-aware continuous modeling offers a practical way to study treatment-response trajectories while preserving individual variation across patients.

When the model was applied to Perturb-seq, it learned perturbation-induced changes in gene expression together with relative changes in cell-state abundance through an unbalanced optimal transport formulation. This formulation enabled the model to calculate cellular growth rates alongside transport plans. Since growth of a cell state due to perturbation implies distinct biological implications to a shift in a single marker value, the growth rate dynamics for gene knockdowns provide an additional population-level summary that complements expression-level predictions. The ability to model both phenotypic shifts and cell-state abundance changes is a notable extension beyond existing balanced flow-based single-cell methods.

The CyTOF signaling dataset tested whether the model can operate on time-resolved data and learn continuous within-condition trajectories from discrete measurements of phosphoprotein dynamics. The coherent predicted signaling responses across *PIK3CA* genotypes imply the model captures significant biological differences in the dose-dependent activation that separates the wild-type from the oncogenic conditions. The correct interpolation of withheld 20 min condition distributions reaffirms the predictive ability of GeoSinkhorn Flow ‘s joint learned latent space and conditional flow matching framework, while connecting discrete condition–time points into a dynamic signaling phenoscape. Furthermore, these results suggest GeoSinkhorn Flow could be applied to pharmacodynamic experiments where continuous flow interpolation could provide greater temporal coverage than experimental assays alone. Despite promising results, the method has some key limitations. The geometric autoencoder is trained on a particular dataset, which means that significant differences between training data and inferred conditions, such as emergent synergistic or antagonistic effects of drug combinations, may reduce the quality of the predictions from the latent geometry. Likewise, the accuracy of the heat kernel approximation is sensitive to graph construction and diffusion parameters, which could produce suboptimal manifold structure if they are not tuned appropriately. When measured time is not available, the inferred progression based on MELD likelihood is also an approximation of the underlying dynamics rather than a direct observation of temporal order. More broadly, the inferred trajectories are model-based reconstructions from unpaired populations rather than directly observed single-cell fates.

Together, these results establish GeoSinkhorn Flow as a principled framework for converting static phenoscapes into predictive dynamic phenoscapes. By unifying manifold geometry learning with conditional flow matching and unbalanced transport, it connects condition-level population states through continuous, geometry-consistent trajectories, explains these transitions through cell-level paths and changes in population mass, and generalizes them to unseen cells, perturbations and intermediate time points. This provides a foundation for understanding how biological systems navigate a phenoscape in response to perturbation.

Future work could extend GeoSinkhorn Flow to integrate multiple molecular modalities and richer representations of experimental conditions, enabling shared dynamic phenoscapes across transcriptomic and proteomic measurements and more complex perturbation spaces.

## 4 Methods

GeoSinkhorn Flow is a novel method that converts a static phenoscape into a dynamic one by linking population-level points with continuous, geometry-consistent trajectories. Existing approaches either treat condition-level distributions as static points or learn only cell-level dynamics. In contrast, GeoSinkhorn Flow learns condition-specific flows between population states.

We first provide an overview of the full algorithm, then describe, in order: the heat-diffusion geometry underlying manifold-aware transport costs; the geometric autoencoder that transfers this geometry into a continuous latent space; progression state construction from measured time or MELD-inferred response bins; Geodesic Sinkhorn couplings between neighboring population states; conditional flow matching that turns static couplings into a predictive vector field; and an optional growth-rate model with unbalanced transport for changes in cell-state abundance.

### 4.1 GeoSinkhorn Flow algorithm

GeoSinkhorn Flow takes as input a collection of unpaired single-cell measurements organized by experimental condition. Each cell is represented by a feature vector *x*_*i*_ ∈ ℝ^*d*^, corresponding to marker intensities or gene-expression measurements. Cells are grouped by condition *l* and progression state *s*, where *s* denotes either a measured experimental time point or a MELD-inferred response bin. We write *µ*_*l,s*_ for the empirical distribution of cells from condition *l* at progression state *s*. The model produces, as output, a learned vector field that can transport any source cell population to its predicted state under a specified condition at any point along a treatment-response trajectory. The first step is to estimate the geometry underlying the data: a weighted nearest-neighbor graph is constructed on the observed cells of the conditions, and heat diffusion on this graph defines geometry-aware distances that follow the densely sampled manifold traced by real cellular states, rather than straight-line separation in high-dimensional feature space. These diffusion distances serve as the ground cost for all subsequent optimal transport computations (Section 4.2).

Because these graph-based diffusion distances are defined only among the conditions used to build the graph, they do not naturally extend to new observations. A geometric autoencoder with encoder *ϕ* : ℝ^*d*^ → ℝ^*k*^ and decoder *ψ* : ℝ^*k*^ → ℝ^*d*^ is therefore trained so that Euclidean distances in its latent space approximate the graph-derived diffusion distances, while the decoder retains the ability to reconstruct marker or gene-expression measurements. Once trained, the encoder maps any new conditions, including held-out or unseen perturbation conditions, into a geometry-aligned latent space in which standard Euclidean optimal transport serves as a tractable proxy for geodesic transport (Section 4.3).

With this geometry-aligned latent space in place, cells must be ordered along a treatment-response axis so that transport can be computed between neighboring population states. When measured time points are available they directly define this ordering; when only cross-sectional snapshots are available, MELD likelihood scores are partitioned into ordered MELD-inferred response bins that stand in for measured time (Section 4.4). Geodesic Sinkhorn optimal transport is then computed between each pair of neighboring states in the sequence, yielding local source–target correspondences that respect the manifold structure of the data (Section 4.5).

These local couplings then supervise a conditional vector field *f*_*θ*_(*z, t, c*_*l*_), where *z* = *ϕ*(*x*) is the encoded cell state and *c*_*l*_ is a condition embedding (encoding drug identity, perturbation gene, or genotype–ligand combination). It takes as input a latent cell state, a condition embedding, and a progression coordinate *t*, and outputs the instantaneous velocity moving that cell toward its coupled target state. Training with a flow-matching objective turns these static, pairwise correspondences into a continuous dynamical model that can generate complete trajectories, interpolate intermediate states, and generalize to conditions not seen during training (Section 4.6). An optional growth-rate model, trained jointly with unbalanced optimal transport, additionally learns which cell states expand or deplete under a given perturbation (Section 4.7).

At inference, new cells are encoded, evolved under the learned condition-specific flow, and decoded back to marker or gene-expression space. Let Ψ_*c,t*_ denote the latent flow induced by *f*_*θ*_ under condition *c*. For an input cell distribution *µ*, the transported distribution is

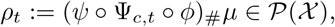

where (·)_#_ denotes the pushforward through the composed encode-flow-decode map. The full algorithm is given in Algorithm 1.

#### Algorithm 1

Training GeoSinkhorn Flow

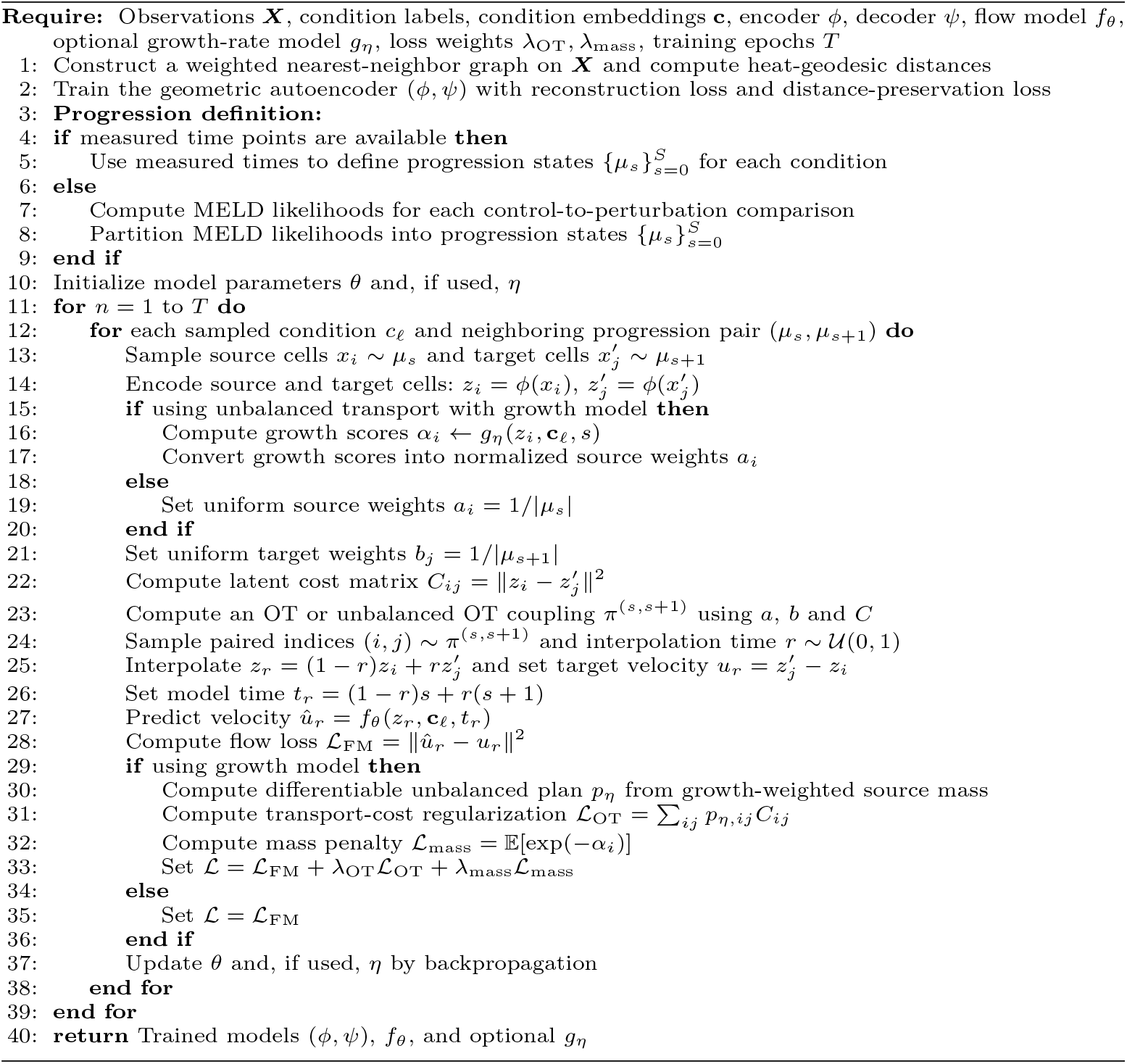

### 4.2 Heat-kernel graph geometry

Geodesic Sinkhorn replaces ambient Euclidean costs with costs derived from diffusion on a graph [2, 9]. Given a weighted nearest-neighbor graph on observed cells, let ***A*** be the adjacency matrix, ***D*** the degree matrix and ***L*** = ***D*** − ***A*** the graph Laplacian. The heat operator is

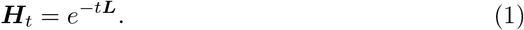

For small diffusion time *t*, heat propagation approximates intrinsic manifold geometry through Varadhan’s formula [10]. This motivates heat-geodesic costs of the form

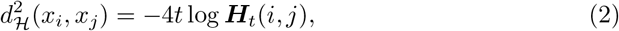

up to implementation-specific normalization. In practice, Geodesic Sinkhorn applies ***H***_*t*_ through sparse graph operations rather than forming a dense eigendecomposition; Chebyshev polynomial approximations are one efficient way to apply this heat operator [11, 12].The following section describes how these graph-derived diffusion distances are transferred into a continuous latent space, so that geometry-aware transport can be extended to unseen cells.

### 4.3 Extending geodesic transport to unseen cells with a geometric autoencoder

The heat-diffusion distances defined in Section 4.2 are computed on a graph built from the observed cells. They do not naturally extend to new observations not present when the graph was constructed. To obtain a continuous representation that generalizes beyond the observed point cloud and that underpins all subsequent transport and flow-matching computations, we train a geometric autoencoder whose latent space preserves the graph-based diffusion geometry.

Let *ϕ* : ℝ^*d*^ → ℝ^*k*^ denote the encoder and *ψ* : ℝ^*k*^ → ℝ^*d*^ the decoder. From the observed cells 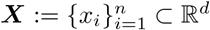, we construct a weighted *k*-nearest-neighbor graph and use the heat kernel operator ***H***_*t*_ to define a diffusion-based distance between observed cell states. The operator ***H***_*t*_ solves the heat equation

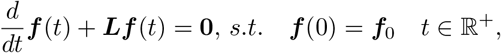

where ***L*** is the graph Laplacian. Inspired by Varadhan’s asymptotic result [13], we define

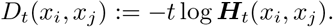

This distance has previously been used in both optimal transport and manifold learning [2, 3, 8, 9]. The autoencoder is trained to reconstruct the original measurements while regularizing latent distances to match *D*_*t*_. We use the geometry-preserving loss

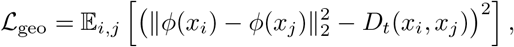

combined with a reconstruction loss on *ψ*(*ϕ*(*x*_*i*_)). The full geometric autoencoder objective is

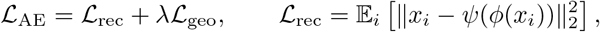

where *λ* controls the strength of the geometry-preserving regularization. This encourages Euclidean distances between latent codes to approximate the diffusion geometry of the observed graph while preserving the measured feature space for downstream interpretation.

After training, a new cell can be mapped to *z* = *ϕ*(*x*) without rebuilding the graph. Optimal transport and flow matching can then be performed using Euclidean costs in this geometry-aligned latent space, and generated latent trajectories can be decoded through *ψ* to recover marker or gene-expression values. The geometric autoencoder thereby transfers the graph-based diffusion geometry into a continuous latent space within which Geodesic Sinkhorn transport and conditional flow matching are subsequently performed. A conditional geometric interpretation of latent straight-line trajectories is provided in Appendix A.3.

### 4.4 Progression state construction

To train a dynamic model from snapshot data, the observed cell populations must be ordered along a treatment-response axis so that transport can be computed between neighboring states. GeoSinkhorn Flow accommodates two distinct situations: datasets in which temporal measurements are directly available, and cross-sectional datasets in which an ordering must be inferred.

For datasets with experimentally measured time, such as the CyTOF signaling data, the observed time points directly define an ordered sequence of progression states. Cells from the same genotype-stimulation condition are grouped by time, and neighboring measured time points are used as source and target populations for Geodesic Sinkhorn transport and flow matching.

For cross-sectional perturbation datasets without measured intermediate time points, such as the PDO drug-response and Perturb-seq experiments, we infer a progression ordering using MELD [5]. MELD is a graph-based method that estimates a continuous experimental signal by smoothing condition labels over the data manifold. Given a control population and a perturbed population, MELD assigns each cell a treatment likelihood score reflecting its position along the control-to-treatment continuum. We compute MELD scores separately for each control-to-perturbation comparison, rank cells by the resulting likelihood, and partition them into a fixed number *S* of ordered bins. We term these *MELD-inferred response bins* to distinguish them from pseudotime coordinates derived from RNA velocity or diffusion pseudo-time: they are not estimates of developmental or temporal order, but estimates of how far each cell has progressed toward the perturbed phenotype. These bins define the empirical distributions 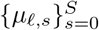 used as input to neighboring-state transport.

In both cases the resulting ordered sequence of population states provides a common interface for the transport and flow-matching stages: measured time points and MELD-inferred response bins are represented identically as progression indices, allowing the same geometry-aware pipeline to be applied to time-resolved and cross-sectional data alike.

### 4.5 Static geometry-aware transport with Geodesic Sinkhorn

Geodesic Sinkhorn provides the static transport component of GeoSinkhorn Flow. The role of this component is to produce, for each pair of neighboring progression states, a coupling matrix that specifies which source cells are matched to which target cells in a way that respects the diffusion geometry of the data manifold. These couplings serve as supervision for the dynamic model described in Section 4.6.

Given two empirical distributions *µ* and *v* supported on observed cells, standardoptimal transport seeks a coupling *π* ∈ Π(*µ, v*) that minimises the total cost oftransporting mass from *µ* to *v*:

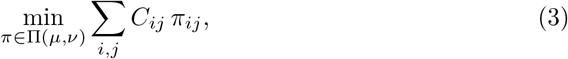

where *C*_*ij*_ is the ground cost between source cell *x*_*i*_ and target cell *x*_*j*_. When this cost is Euclidean, transport can create shortcuts across sparsely sampled or biologically unsupported regions of a nonlinear cellular manifold. Geodesic Sinkhorn avoids this by defining *C*_*ij*_ from the heat-diffusion distances, so that transport is encouraged to follow densely sampled cellular trajectories rather than straight lines in ambient feature space. The entropy-regularised version of this problem is solved efficiently with Sinkhorn iterations, yielding a geometry-aware coupling *π*^*^ between the two populations. Full notation for entropy-regularised optimal transport is provided in Appendix A.1.

In GeoSinkhorn Flow, Geodesic Sinkhorn couplings are computed between each pair of neighboring progression states (*µ*_*l,s*_, *µ*_*l,s*+1_) for each condition *l*. These local couplings respect the manifold geometry and provide the source-target correspondences that supervise the conditional flow model. The graph-based construction, however, is tied to the observed point cloud: it defines distances and couplings only among measured cells, and does not by itself yield a continuous representation for unseen cells or a vector field for generating trajectories. hese local couplings respect the manifold geometry and provide the source–target correspondences that supervise the conditional flow model.

### 4.6 Learning continuous dynamics with conditional flow matching

The Geodesic Sinkhorn couplings computed in the preceding section are static: for each pair of neighboring progression states, they specify which source cells correspond to which target cells, but they do not define a trajectory between them or a mechanism for generating responses at intermediate states or under unseen conditions. Conditional flow matching [14, 15] converts these local, pairwise correspondences into a continuous, condition-aware vector field. OT-coupled flow-matching notation is summarised in Appendix A.2.

For a condition *l* and neighboring progression states (*µ*_*l,s*_, *µ*_*l,s*+1_), we encode source and target cells into the geometry-aligned latent space. Let *z*_0_ = *ϕ*(*x*_0_) and *z*_1_ = *ϕ*(*x*_1_) denote a source-target pair sampled from the geometry-aware coupling between these neighboring states. We define a linear interpolation between the paired latent states,

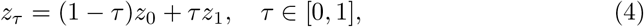

with target velocity *u*_*τ*_ = *z*_1_−*z*_0_. The interpolation variable *τ* describes local movement between two adjacent progression states; the global progression coordinate is obtained by interpolating between the neighboring state indices, *t*_*τ*_ = (1 − *τ* ) *s* + *τ* (*s* + 1).

The conditional flow model is a vector field *f*_*θ*_ : ℝ^*k*^ × ℝ^*c*^ × [0, 1] → ℝ^*k*^, which takes as input the current latent state, a condition embedding *c*_*l*_, and the global progression coordinate, and predicts the instantaneous velocity in latent space. We train *f*_*θ*_ with the flow-matching objective

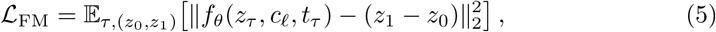

where (*z*_0_, *z*_1_) is sampled according to the transport coupling for the corresponding neighboring population pair. In contrast to standard flow matching in ambient measurement space, this objective is trained in a latent space whose Euclidean geometry approximates the diffusion geometry of the observed phenoscape, a property enforced by the geometric autoencoder. The condition embedding *c*_*l*_ allows the same initial cell state to evolve toward different endpoints under different experimental conditions.

At inference, a new cell is encoded into latent space and evolved under the learned ordinary differential equation d*z*_*t*_*/*d*t* = *f*_*θ*_(*z*_*t*_, *c*_*l*_, *t*) for a specified condition embedding and progression interval. The resulting latent trajectory is decoded through *ψ* to obtain a continuous, condition-aware trajectory in the original marker or gene-expression space. Conditional flow matching thereby turns local, static Geodesic Sinkhorn correspondences into a vector field that can generate responses at arbitrary intermediate progression values and for new input cells. A consistency statement for the learned trajectories is provided in Appendix A.4.

### 4.7 Growth Rate Model and Unbalanced Optimal Transport

The balanced transport framework assumes that the total mass of cells is conserved across neighboring progression states, every source cell is transported to exactly one target cell. This assumption is appropriate when perturbations shift cell states without changing the relative abundance of those states. When a perturbation also causes certain cell states to expand or deplete, balanced transport will misattribute abundance changes to state transitions. To model both effects simultaneously, we extend the framework with a learnable growth-rate model and an unbalanced transport objective.

Let *g*_*η*_ : ℝ^*k*^ × ℝ^*c*^ × [0, 1] → ℝ be a neural network that takes a latent cell state *z*, a condition embedding *c*_*l*_, and a progression value *t*, and returns a scalar score for the relative mass contribution of that cell under the given condition. For a source population 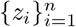, these scores define normalised source weights

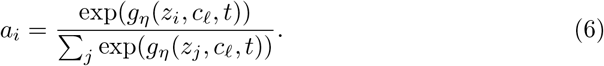

A transport coupling is then computed between source weights *a* = (*a*_*i*_) and the target population using the latent-space cost matrix 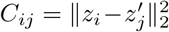 . When the unbalanced variant is enabled, this coupling is computed with entropic regularisation *ε* and marginal relaxation *ρ*, so that transported source and target marginals are penalised rather than enforced exactly. This mass-aware coupling is then used to sample training pairs for the conditional flow model, so that the learned dynamics account for both state transitions and changes in the relative abundance of cell states.

The total training objective combines the flow-matching loss with a transport-cost regularisation term induced by the growth-weighted unbalanced transport plan:

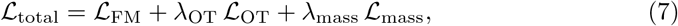

where 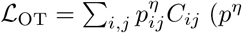 being the differentiable unbalanced transport plan computed from growth-weighted source mass) and *L*_mass_ = E_*i*_[exp(−*g*_*η*_(*z*_*i*_, *c*_*l*_, *t*))]. The growth-rate model *g*_*η*_ is used only when the unbalanced transport variant is enabled; the balanced variant sets all source weights uniformly and optimizes only L_FM_.

### 4.8 Data Sources and Preprocessing

#### 4.8.1 Synthetic datasets

We generated three synthetic datasets to evaluate geometry-aware distance recovery under controlled manifold structure: a Swiss roll, a saddle surface and a toroidal surface. Each dataset consisted of 100 local distributions, with 50 sampled points per distribution, yielding 5,000 points per manifold. The distribution centers were sampled over the corresponding intrinsic coordinates of each manifold, and local noise was added around each center to create finite-width populations rather than isolated points. For each dataset, we computed ground-truth manifold distances from the known generating coordinates or analytic distance approximations. These distances were used to evaluate whether Euclidean distances in the learned autoencoder latent space recover the geometry-aware distance structure of the original manifold. Detailed generation procedures and distance calculations are provided in Appendix B.2.

#### 4.8.2 PDO dataset

The PDO dataset was obtained from the colorectal cancer patient-derived organoid drug-screening study of Ramos Zapatero et al. [4]. The source dataset comprises 10 colorectal cancer patient-derived organoid (PDO) lines profiled by single-cell mass cytometry in both monoculture and co-culture with cancer-associated fibroblasts (CAFs). Using TOBis (thiol-reactive organoid barcoding in situ) for high-throughput multiplexing, the study measured 44 antibodies spanning protein and post-translational modification signaling, DNA damage, apoptosis and cell state, generating a highly multiplexed collection of more than 2,500 PDO/CAF cultures across 10 patients. For the analyses presented here, we restricted attention to monoculture PDOs so that the learned trajectories reflect drug-induced cancer-cell state changes without the additional variability introduced by stromal co-culture.

The source screen included the chemotherapies SN-38 (S), 5-fluorouracil (5-FU) (F) and oxaliplatin (O), together with the targeted agents cetuximab (C), VX-970 (berzosertib) (V) and LGK-974 (L). In our analysis, each perturbation was compared with its matched control condition from the same patient and culture setting, and the treatment set included both single and combination conditions. Single-treatment conditions were used for training, whereas combination-treatment conditions were withheld for evaluation of compositional generalization.

For preprocessing, we retained the per-cell marker profiles provided in the source dataset and organized cells by patient, control and treatment condition. MELD scores were computed separately for each control-to-treatment comparison and used to define pseudotime bins that order cells along a treatment-response continuum. These pseudotime bins were then used to train the GeoSinkhorn Flow model.

#### 4.8.3 Perturb-seq dataset

The Perturb-seq dataset was obtained from the coronary artery disease CRISPR interference screen in human arterial endothelial cells reported by Schnitzler et al. [6]. Candidate CAD-associated genes were knocked down using CRISPR interference, and the resulting transcriptional states were measured by single-cell RNA sequencing. We constructed the response model using 5,000 highly variable genes after batch correction. Negative-control cells were retained as the source population for control-to-perturbation transport.

For the main analysis, we used a biologically motivated subset of perturbations spanning endothelial CAD programs highlighted in the original study, including CCM signalling, flow/KLF response, angiogenesis, endothelial migration, extracellular matrix remodelling and inflammatory activation. Perturbations were required to have sufficient cell counts and an available gene embedding. Six CAD-relevant perturbations were withheld from conditional flow training to evaluate generalization to unseen perturbation genes: *CCM2, MAP2K5, EDN1, NOS3, TSPAN14* and *PECAM1*. The remaining selected non-control perturbations were used for training. Held-out perturbation cells were excluded from model training, whereas negative-control cells were retained in both training and evaluation to provide a shared source distribution.

Because the Perturb-seq screen contains endpoint measurements rather than measured intermediate time points, progression states were inferred separately for each control-to-perturbation comparison. MELD was applied to the corresponding negative-control and perturbed cells using the first 10 principal components. Treatment likelihood scores were normalized and partitioned into five pseudotime bins. Neighboring bins were then used as source and target populations for geometry-aware transport and conditional flow matching. Details of cell-count filtering and perturbation selection are provided in Appendix B.4.

##### Gene embedding

For the Perturb-seq experiments, perturbation identity was represented using fixed gene embeddings generated with GeneWalk [16]. GeneWalk uses random-walk-based network representation learning to embed genes in a functional graph defined by gene annotations and Gene Ontology relationships. We generated embeddings from the Ensembl gene identifiers present in the Perturb-seq expression matrix, using GeneWalk with Ensembl identifiers as input and a fixed random seed. The resulting node embeddings were eight-dimensional, and the gene-level vectors were used as perturbation-conditioning vectors in GeoSinkhorn Flow .

Perturbation labels were matched to GeneWalk embeddings by gene symbol. Symbols were converted to upper case before lookup, and obsolete or alternative symbols were resolved using MyGene.info when possible. Labels with the suffix `-TSS2` were mapped to the corresponding gene symbol, and composite labels containing `-AND-` were represented by the mean of the available component gene embeddings. Negative-control cells were assigned an all-zero condition vector. Perturbations without an available GeneWalk embedding were removed from embedding-conditioned training and evaluation.

The gene embedding was used as the condition vector in GeoSinkhorn Flow . Each training batch contained cells from a single perturbation, and the corresponding GeneWalk vector was provided to the conditional vector field together with the latent cell state and the progression-time embedding. Thus, the GeneWalk embeddings supplied a fixed functional representation of perturbation identity, while the flow-matching model learned perturbation-specific cellular dynamics from the observed Perturb-seq response distributions.

#### 4.8.4 CyTOF signaling dataset

The CyTOF signaling dataset was obtained from the HeLa spheroid CyTOF experiments reported by Madsen et al. [7]. We downloaded the single-cell files from the authors’ OSF repository (https://osf.io/nztm6/files/m36vt). The dataset profiles cycling, non-apoptotic HeLa spheroid cells with wild-type *PIK3CA* or one (1×H1047R) or two (2×H1047R) endogenous copies of *PIK3CA*^H1047R^ following stimulation with EGF or IGF1 after 4 h of serum starvation. Although the source study also includes BYL719 perturbations, the analyses shown in Figure 5 were restricted to the H_2_O vehicle-control conditions under EGF and IGF1.

For our analysis, marker intensities were transformed using the inverse hyperbolic sine transform with cofactor 5, *x* 1→ arcsinh(*x/*5), matching the normalization used in the source study. We modeled eight signaling markers summarizing the major pathway responses in this experiment: pAKT^S473^, pERK1/2^T202*/*Y204^, pS6^S240*/*S244^, pNDRG1^T346^, pSMAD2/3, total S6, pRB^S807*/*S811^ and cleaved caspase-3^D175^. For the EMD-PHATE analysis, we used the subset of five signaling markers emphasized in the original CyTOF study: pAKT^S473^, pERK1/2^T202*/*Y204^, pS6^S240*/*S244^, pNDRG1^T346^ and pSMAD2/3. To assess temporal generalization, we withheld the 20 min time point for evaluation and used the remaining time points to train the GeoSinkhorn Flow .

## Supplementary information

If your article has accompanying supplementary files please state so here.

Authors reporting data from electrophoretic gels and blots should supply the full unprocessed scans for key as part of their Supplementary information. This may be requested by the editorial team/s if it is missing.

Please refer to Journal-level guidance for any specific requirements.

## Acknowledgements

Acknowledgements are not compulsory. Where included they should be brief. Grant or contribution numbers may be acknowledged.

Please refer to Journal-level guidance for any specific requirements.

## Declarations

## Appendix A Computational and theoretical background

This appendix collects the computational notation and theoretical results that support the GeoSinkhorn Flow methodology.

### A.1 Entropy-regularized optimal transport

Let *µ* and *v* be empirical distributions supported on source cells 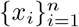 and target cells 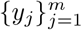 . A transport plan 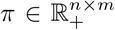 assigns mass from source to target while matching the prescribed marginals. For a ground-cost matrix *C*_*ij*_, entropy-regularized OT solves

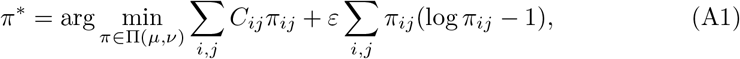

where *ε >* 0 controls the strength of entropic smoothing [17, 18]. In GeoSinkhorn Flow, the coupling *π*^*^ is computed between neighboring progression states and then used to sample source–target training pairs for the flow model. When growth or transition weights are used, the source marginal is learned rather than fixed uniformly, and marginal relaxation penalizes rather than strictly enforces deviations from the prescribed marginals, giving the marginally relaxed mass-aware variant described in Methods.

### A.2 OT-coupled conditional flow matching

Dynamic OT represents transport through a time-indexed density *ρ*_*t*_ and vector field *v*_*t*_ satisfying the continuity equation, rather than only through a static endpoint coupling [19]. Flow matching learns such a vector field by regressing velocities along sampled probability paths [14, 15]. For a coupled latent pair (*z*_0_, *z*_1_) sampled from *π*^*^, GeoSinkhorn Flow uses the standard linear conditional path

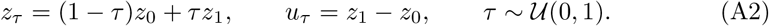

The conditional vector field *f*_*θ*_(*z*_*τ*_, *c, t*_*τ*_ ) is trained with

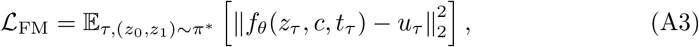

where *c* is the perturbation or treatment embedding and *t*_*τ*_ is the local progression coordinate. The role of the OT coupling is to choose biologically plausible source– target pairs before fitting the continuous vector field.

#### A.3 Geometric interpretation (conditional)

This section gives a geometric reading of the latent-space straight-line CFM trajectory. The result is conditional on a verifiable geometric assumption about the encoder and is presented as such, not as a guaranteed property.

##### Assumption 1

(Local isometric encoder) There exists a Riemannian manifold (ℳ, *g*) and an open set *U* ⊂ ℳ on which the data lie, together with a smooth map Φ : *U* → ℝ^*k*^ that is a diffeomorphism onto its image and a local isometry: for every *x* ∈ *U* and *u, w* ∈ *T*_*x*_ℳ,

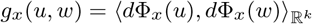

##### Theorem 2

(Latent straight lines pull back to manifold geodesics) *Under Assumption 1, let γ* : [0, 1] → Φ(*U* ) *be a Euclidean straight line, γ*(*t*) = (1 − *t*)*y*_0_ + *ty*_1_. *Then the pullback curve*

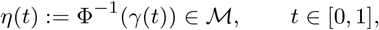

*is a length-minimizing geodesic on* (ℳ, *g*) *joining* Φ^−1^(*y*_0_) *and* Φ^−1^(*y*_1_), *provided the entire image γ*([0, 1]) *lies in* Φ(*U* ).

*Proof* The arc-length of any smooth curve *β* : [0, 1] → *U* on (ℳ, *g*) is

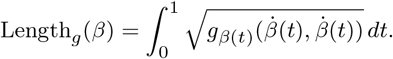

By the local isometry,

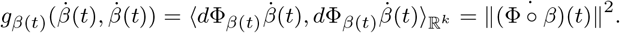

Hence Length_*g*_ 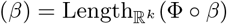 .

Now let *η*(*t*) = Φ^−1^(*γ*(*t*)). Then Φ° *η* = *γ* is a Euclidean straight line, which is the unique length-minimizing curve from *y*_0_ to *y*_1_ in ℝ^*k*^. Therefore *η* is the unique length-minimizer in (ℳ, *g*) from Φ^−1^(*y*_0_) to Φ^−1^(*y*_1_), hence a geodesic. □

##### Remark 1

(Empirical verifiability) Assumption 1 is strong but verifiable. PHATE embeddings [8], under appropriate diffusion-time and bandwidth choices, approximate isometries on the diffusion-geometry manifold. The assumption can be empirically tested: for sample pairs (*x*_1_, *x*_2_) ∈ *U* × *U* with known geodesic distance, check that

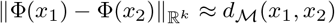

within a tolerance that depends on the local curvature scale. For our pipeline, deviations from exact isometry contribute additional error but do not invalidate the qualitative conclusion that latent straight-line trajectories track manifold geodesics.

### A.4 Consistency of learned trajectories

We now show that the simulated trajectory generated by the learned vector field *v*_*ϕ*_ converges to the true population dynamics in *W*_2_, with a rate depending on the initial sampling error and the within-bin training error.

#### Theorem 3

(ODE flow stability under velocity perturbation) *Let v*^*^, *v*_*ϕ*_ : ℝ^*d*^ × [0, 1] → ℝ^*d*^ *be two velocity fields with v*^*^ *uniformly L-Lipschitz in x on* [0, 1]. *Let X*_0_ ∼ *µ*_0_ *be a random initial condition, and define*

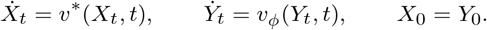

*Define the worst-case trajectory error*

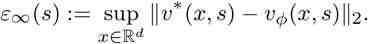

*Then for every t* ∈ [0, 1], *almost surely*,

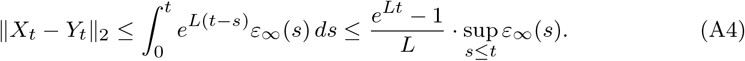

*In particular, W* (Law(*X* ), Law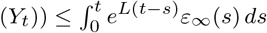

*Proof* Set *Z*_*t*_ := *X*_*t*_ − *Y*_*t*_, *Z*_0_ = 0. Then

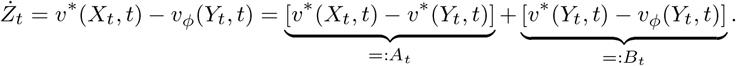

By Lipschitzness of *v*^*^, ∥*A*_*t*_∥ ≤ *L*∥*Z*_*t*_∥. By definition, ∥*B*_*t*_∥ ≤ *ε*_∞_(*t*). Therefore

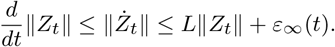

Gronwall’s inequality yields

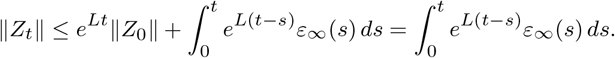

Bounding *ε*_∞_(*s*) ≤ sup_*s*≤*t*_ *ε*_∞_(*s*) gives the second inequality. The Wasserstein bound follows from the synchronous coupling (*X*_*t*_, *Y*_*t*_). □

We now apply this to the bin-trained CFM setting. Let 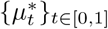 be the true population law in *P*_2_(ℝ^*d*^), with associated true velocity *v*^*^ satisfying 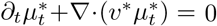

#### Assumption 4

(Regularity of true dynamics) 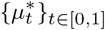 is absolutely continuous in (*P*_2_(ℝ^*d*^), *W*_2_), with metric derivative 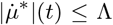 for a.e. *t*, and *v*^*^ is uniformly *L*-Lipschitz in *x* on [0, 1].

#### Assumption 5

(Bin-pair training error) For each *k* = 1, …, *K*, the within-bin trained vector field 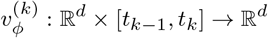 satisfies

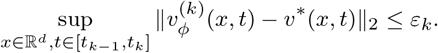

The piecewise field 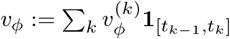 inherits uniform error *ε* := max_*k*_

#### Theorem 6

(Discrete-to-continuous consistency) *Under Assumptions 4 and 5, let X*_*t*_ *solve* 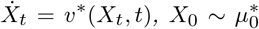, *and let Y*_*t*_ *solve* 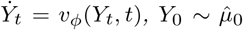 *with* 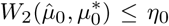 . *Then*

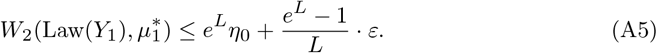

*In particular, refining bins and increasing initial sample size drives W*_2_(Law(*Y*_1_), 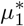 ) → 0 *when the within-bin error ε and initial sampling error η*_0_ *vanish*.

*Proof* Couple *X*_0_ and *Y*_0_ optimally so that 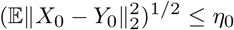. Applying Theorem 3 to the coupled trajectories with initial offset *Z*_0_ = *X*_0_ − *Y*_0_ gives

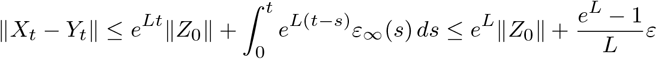

for *t* = 1. Taking *L*^2^ expectation and using the optimal initial coupling yields

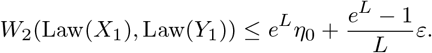

Since Law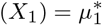, this concludes. □

#### Remark 2

(Bin-width controls *ε*_*k*_) The connection between bin width *δ*_*K*_ and within-bin training error *ε*_*k*_ follows standard CFM consistency intuition [14, 15]. Under additional smoothness, the OT-displacement velocity from 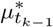 to 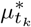 approximates 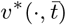 on [*t*_*k*−1_, *t*_*k*_], and the population CFM minimizer matches this displacement velocity. Consequently, *ε*_*k*_ decreases as bins become finer, up to finite-sample and stochastic optimization error.

#### Corollary 7

(Forward monotonicity preservation) *Under Assumption 4 and additionally that v*^*^ *is the gradient of a progression potential ψ*^*^, *if ε is small enough that* 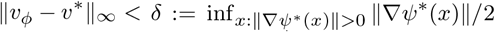, *then the simulated trajectory Y*_*t*_ *satisfies* 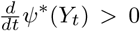 *wherever* ∇*ψ*^*^(*Y*_*t*_) ≠0.

*Proof* Since *v*^*^ = ∇*ψ*^*^,

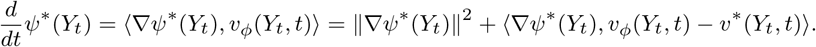

Thus

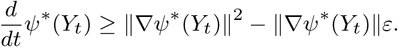

Whenever ∥∇*ψ*^*^(*Y*_*t*_)∥ *>* 2*ε*, the bound is positive. □

## Appendix B Additional experimental details and results

### A.1 Implementation details and hyperparameters

GeoSinkhorn Flow was implemented in Python using PyTorch [20], PyTorch Lightning [21], Hydra [22], POT [23], and TorchCFM [24]. Unless otherwise stated, experiments used a fixed random seed of 42. Model configuration, checkpointing, logging and dataset-specific overrides were managed with Hydra configuration files.

Models were trained in two stages. First, we learned a geometric autoencoder representation of the single-cell state space. The geometric autoencoder used multilayer perceptron encoder and decoder networks with ReLU activations and was trained with the reconstruction and geometry-preserving losses described in Methods. Architecture and optimization settings were selected separately for each dataset and are summarized in Table B1. Unless otherwise specified, geometric autoencoders were optimized with AdamW, Gaussian input noise was added during training, gradients were clipped at 1.0 and early stopping was performed on validation loss.

Second, we trained a condition-aware flow-matching model in the learned latent space. The vector field was parameterized by a conditional multilayer perceptron that received the latent state, condition embedding and a Gaussian Fourier embedding of the progression variable as inputs. Training pairs were sampled from optimal transport couplings between source and target mini-batches. Flow-matching and optimal-transport hyperparameters are summarized in Table B1.

For the PDO and Perturb-seq experiments, which did not include measured intermediate time points, cells were assigned to five MELD-based progression bins. MELD was run with *k* = 15 and *β* = 67 for both datasets. The input representation differed by dataset: PDO used all measured markers, whereas Perturb-seq used the first 10 principal components.

A small neural network predicted positive source-cell mass weights for growth-aware unbalanced transport using a Softplus output. The growth model used hidden layers of sizes 64 and 32, ReLU activations and progression time as an additional covariate.

**Table B1.** Dataset-specific hyperparameters used in GeoSinkhorn Flow experiments. Dashes indicate settings that were not applicable or were not used for the corresponding dataset.

| Hyperparameter | Synthetic | PDO | CyTOF signaling | Perturb-seq |
| --- | --- | --- | --- | --- |
| AE width | 256 | 64 | 512 | 128 |
| AE latent dimension | 512 | 64 | 512 | 128 |
| AE batch size | 128 | 256 | 128 | 128 |
| AE learning rate | $10^{-4}$ | $10^{-3}$ | $10^{-3}$ | $10^{-3}$ |
| AE regularization $\lambda$ | 10 | 10 | 10 | 10 |
| Heat-geodesic $k$ | 5 | 5 | 5 | 5 |
| Heat time $\tau$ | 5.0 | 5.0 | 5.0 | 5.0 |
| Flow model batch size | – | 256 | 128 | 32 |
| Flow model width | – | 256 | 512 | 1024 |
| Flow model learning rate | – | $10^{-3}$ | $10^{-3}$ | $10^{-3}$ |
| OT entropy regularization $\varepsilon$ | – | 0.1 | 0.1 | 0.1 |
| OT marginal relaxation $\rho$ | – | 1.0 | 1.0 | 2.0 |
| Flow interpolation noise $\sigma$ | – | 1.0 | 0.5 | 1.0 |

### B.2 Synthetic experiments

To test and evaluate the proposed method’s performance on distinct geometric structures and get a detailed understanding of how the method performs on these structures, we constructed three different synthetic datasets: the Swiss Roll, Toroidal Surface, and Saddle Surface datasets. Each dataset aims to capture unique geometric properties, noise characteristics, and distribution patterns.

#### B.2.1 Generation of synthetic datasets

##### Swissroll dataset

The Swiss Roll dataset is constructed to demonstrate a two-dimensional plane rolled into a three-dimensional space. By default, the dataset consisted of 100 distributions, each containing 50 points, resulting in a total of 5000 points. We first generate random means of *t* and *y* and add additional noise to them to create variability around the mean. The (*y, t*) coordinates contain the geometric information of the Swiss Roll Dataset. We then transform the two-dimensional coordinates into three-dimensional coordinates (*x, y, z*) using the following formula: *x* = *t* · *cos*(*t*) and *z* = *t* · *sin*(*t*), while *y* retained its noisy values. Optionally, the dataset could be rotated to a higher dimension using random orthogonal transformations to test the robustness of algorithms to dimensional changes. To calculate the pairwise geodesic distance, we first unroll the *t* by 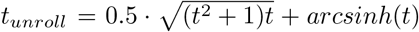, then using the Euclidean distance on the (*y, t*_*unroll*_) space to approximate the geodesic distance.

##### Toroidal surface dataset

The Toroidal Surface dataset simulated a toroidal structure, incorporating periodic boundary conditions. By default, the dataset also consisted of 100 distributions, each containing 50 points, resulting in a total of 5000 points. To begin, we uniformly generated the random mean values for the angular coordinates *θ* and *ϕ* to define the toroidal structure. To introduce the variability around the mean, we added random noise to *θ* and *ϕ*. The noisy two-dimensional coordinates (*θ, ϕ*) were transformed into three-dimensional Cartesian coordinates (*x, y, z*) following the formulation: *x* = (width + height · cos(*θ*)) · cos(*ϕ*), *y* = (width + height · cos(*θ*)) · sin(*ϕ*), and *z* = height · sin(*θ*), where the width and height are the user-defined parameters. Additional noise could be added to the Cartesian coordinates to further simulate the real-world data imperfection. Like the Swiss Roll dataset, the dataset could be rotated to a higher dimension using random orthogonal transformations. Pairwise geodesic distances were approximated by the Euclidean distance, considering periodic boundary conditions on the torus. Specifically, the pairwise geodesic distance is acquired from:

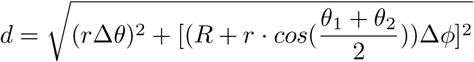

Where r is the radius of the pipe, determining the width of the pipe, and R is the major radius of the torus, determining its overall size of the Toroidal.

##### Saddle surface dataset

The Saddle Surface dataset defines a saddle-shaped surface, which also contains 100 distributions, each having 50 points, resulting in a total of 5000 points. Mean values for the parameters *u* and *v* were sampled uniformly to define the saddle structure. Like the Swiss Roll dataset and the Toroidal dataset, noise was added to these parameters to incorporate the variability within the dataset. The noisy *u* and *v* values were transformed into three-dimensional Cartesian coordinates (*x, y, z*) using the saddle equation *z* = *a* · *u*^2^ − *b* · *v*^2^, with *x* = *u* and *y* = *v*. Pairwise geodesic distances were approximated by the Euclidean distance in the parameters space defined by *u* and *v*, defined by:

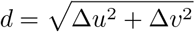

#### B.2.2 Distance comparisons on synthetic manifolds

For each synthetic dataset, we evaluated distances at the level of distributions rather than individual points. Let 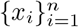 denote the sampled support points on the manifold, and let 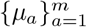 denote the local probability distributions supported on these points. Each method defines a ground cost between support points, which is then used to compute a pairwise Wasserstein distance between distributions. The resulting *m* × *m* distance matrix is compared against a ground-truth distribution-distance matrix.

The ground-truth reference distance is computed using the known geodesic distance on the synthetic manifold. Specifically, we compute the pairwise geodesic distance *d*_geo_(*x*_*i*_, *x*_*j*_) between support points, square it to form the ground cost, and compute Wasserstein distances between all pairs of distributions:

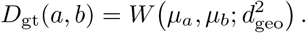

This provides the reference geometry-aware distance between distributions.

We compare several approximations to this reference. The ambient Euclidean baseline uses squared Euclidean distances in the observed coordinate space,

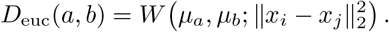

This baseline can be inaccurate on curved or folded manifolds because it may create shortcuts through unsupported regions of the ambient space.

The HeatGeo distance [3] uses a graph constructed from the observed samples to approximate intrinsic geometry. Heat diffusion on this graph defines a geometry-aware ground cost *d*_HeatGeo_(*x*_*i*_, *x*_*j*_), yielding

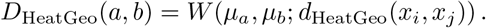

This distance is sample-based and graph-dependent, so it captures the observed manifold geometry but does not directly define distances for new observations.

The learned autoencoder distance evaluates whether the geometric autoencoder transfers this graph-based geometry into a continuous latent representation. Let *f*_*θ*_(*x*) denote the encoder. We compute squared Euclidean distances in latent space,

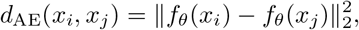

and use them as the ground cost for Wasserstein distances:

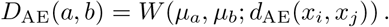

**Table B2.** Quantitative comparison of geometry-aware distance recovery on synthetic manifolds. Pairwise Wasserstein distance matrices between local distributions were computed on the Swiss roll, saddle surface and toroidal surface using each ground-distance method, then compared with the ground-truth geodesic Wasserstein distance matrix. Spearman and Pearson correlations measure global agreement with the reference distances, while *P* @2 and *P* @5 measure nearest-neighbor preservation among distributions. Values are reported as mean ± standard deviation; higher values indicate better recovery of the ground-truth geometry.

| Dataset | Method | Spearman | Pearson | P@2 | P@5 |
| --- | --- | --- | --- | --- | --- |
| Swiss Roll | Model | 0.881 ± 0.017 | **0.826 ± 0.013** | **0.893 ± 0.006** | **0.962 ± 0.009** |
| | GeoSinkhorn | **0.918 ± 0.020** | 0.73 ± 0.009 | 0.538 ± 0.04 | 0.819 ± 0.050 |
| | HeatGeo | 0.909 ± 0.013 | 0.695 ± 0.012 | 0.528 ± 0.046 | 0.802 ± 0.046 |
| | Diffusion EMD | 0.896 ± 0.008 | 0.758 ± 0.014 | 0.525 ± 0.068 | 0.809 ± 0.041 |
| | Euclidean | 0.474 ± 0.037 | 0.424 ± 0.031 | 0.912 ± 0.023 | 0.960 ± 0.011 |
| Saddle | Model | 0.861 ± 0.011 | 0.871 ± 0.007 | **0.675 ± 0.026** | **0.764 ± 0.020** |
| | GeoSinkhorn | 0.933 ± 0.018 | 0.869 ± 0.008 | 0.608 ± 0.018 | 0.720 ± 0.015 |
| | HeatGeo | **0.937 ± 0.015** | 0.836 ± 0.007 | 0.618 ± 0.012 | 0.745 ± 0.011 |
| | Diffusion EMD | 0.935 ± 0.010 | **0.876 ± 0.011** | 0.635 ± 0.010 | 0.749 ± 0.018 |
| | Euclidean | 0.535 ± 0.021 | 0.498 ± 0.021 | 0.317 ± 0.021 | 0.449 ± 0.018 |
| Toroidal | Model | 0.919 ± 0.005 | 0.886 ± 0.004 | 0.73 ± 0.013 | 0.787 ± 0.012 |
| | GeoSinkhorn | 0.903 ± 0.006 | 0.840 ± 0.006 | 0.762 ± 0.026 | 0.837 ± 0.007 |
| | HeatGeo | 0.936 ± 0.002 | 0.899 ± 0.005 | 0.778 ± 0.020 | 0.863 ± 0.001 |
| | Diffusion EMD | **0.946 ± 0.002** | **0.925 ± 0.004** | 0.757 ± 0.013 | 0.857 ± 0.003 |
| | Euclidean | 0.936 ± 0.004 | 0.923 ± 0.006 | **0.937 ± 0.026** | **0.919 ± 0.020** |

Agreement between *D*_AE_ and *D*_gt_ indicates that Euclidean distances in the learned latent space recover the intrinsic distribution-level geometry of the original manifold.

We additionally evaluate graph-based transport distances, including GeoSinkhorn [2] and diffusion EMD [25]. These methods operate directly on the HeatGeo graph rather than through a precomputed dense ground-cost matrix or learned latent embedding. GeoSinkhorn computes entropy-regularized transport using heat-kernel operations on the graph, while diffusion EMD embeds distributions using multiscale diffusion geometry and compares them in the resulting representation.

For each method, we compare the induced distribution-distance matrix to the ground-truth matrix *D*_gt_ using rank and correlation metrics, including Spearman correlation, Pearson correlation, and nearest-neighbor preservation scores. The complete quantitative comparison is summarized in Table B2.

#### B.3 PDO additional details and results

Cell-cycle status was assessed at the single-cell level using marker-intensity gating. Normalized marker intensities were extracted for a panel of cell-cycle and cell-state markers, including pRB, IdU, pHH3, Cyclin B1, cPARP and pHistone H2A. Fibroblast cultures were excluded, and analyses were restricted to the highest treatment concentration for each patient–treatment condition. For each marker, intensity distributions were visualized across treatment groups and thresholds were applied to separate marker-negative and marker-positive populations (Figure B1). These gates were used to compare treatment-associated shifts in cell-cycle and cell-state phenotypes: IdU positivity indicated DNA synthesis and S-phase entry, pHH3 marked mitotic cells, Cyclin B1 reflected G2/M-associated accumulation, pRB reflected cell-cycle regulatory activity, cPARP indicated apoptotic cells and pHistone H2A marked DNA-damage- or stress-associated signaling.

Supplementary Figure B2a shows the generalization of cellular response trajectories to unseen drug combinations through compositional treatment encoding. With this encoding, all the PDOs were embedded into a shared latent space learned by a common geometric autoencoder. This shared representation provides a common coordinate system for comparing treatment-response trajectories across PDOs while still allowing each PDO to retain patient-specific response structure.

**Fig. B1.**
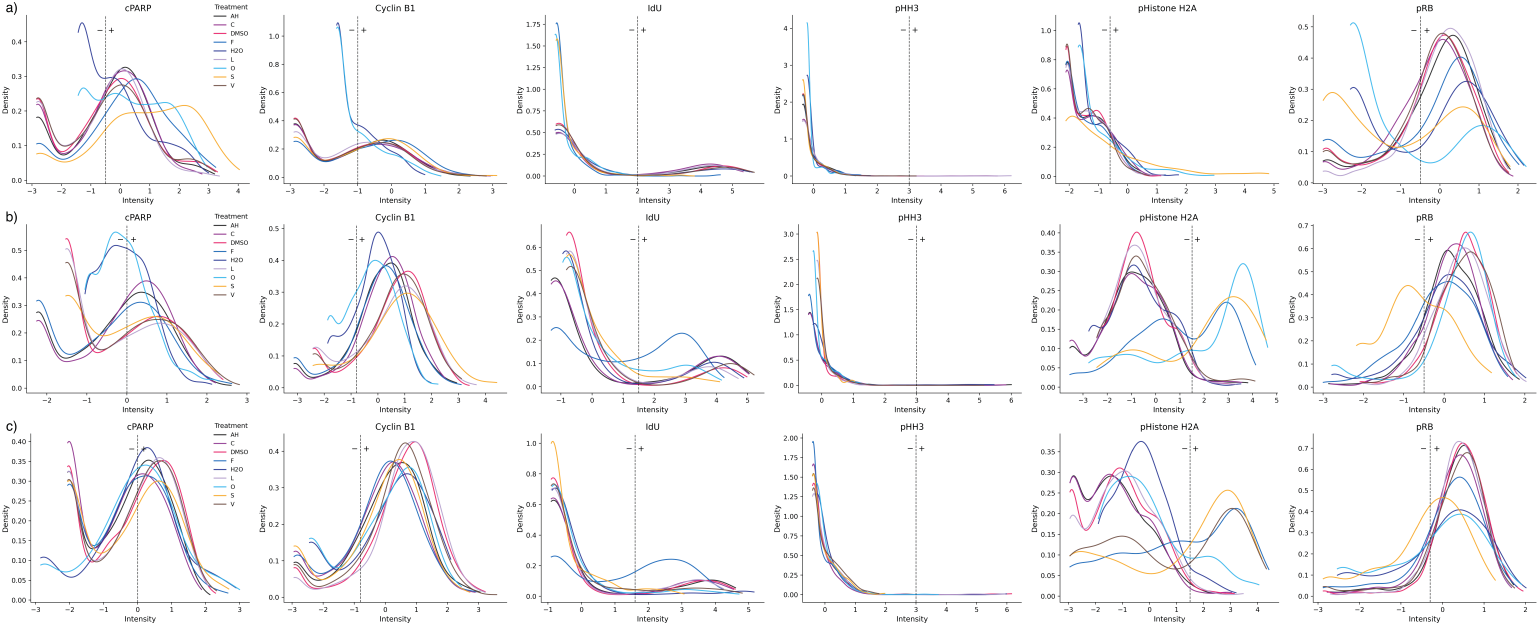
Cell-cycle gating thresholds for cPARP, Cyclin B1, IdU, pHH3, pHistone and pRB for a) PDO 21, b) PDO23 and c) PDO 27. Cell-state thresholds were applied to batch-mean-centered, arcsinh-transformed intensities and were set using marker-specific prior knowledge, with manual refinement at population boundaries following standard cytometry gating conventions.

For PDO21, Figure 3d-g and Supplementary Figure B2b-f summarize the inferred trajectories, marker-level changes, growth-rate landscape and cell-cycle phase assignments. Supplementary Figure B2b shows the MELD-based ordering used to construct the PDO21 trajectories, while Supplementary Figure B2c-f provides additional outputs for Oxaliplatin, growth-rate inference and cell-cycle phase. Together with the main PDO21 panels, these results show that the predicted trajectories move from control-like cell states toward treatment-associated apoptotic states, with marker-level trends consistent with increased cPARP and pHistone H2A and reduced IdU and pRB. We then show the corresponding inferred trajectory, cell-cycle, growth-rate and marker-level outputs for PDO23 and PDO27 in Supplementary Figures B3 and B4. Compared with PDO21, PDO23 and PDO27 exhibit greater heterogeneity in their response to treatment. In both PDO23 and PDO27, SN-38 and 5-FU each increase IdU and pH2AX while decreasing pRB, though 5-FU shows an initial spike in IdU before tapering to a sustained net increase. For Oxaliplatin, treatment response differs between PDO23 and 27: PDO23 shows rising pH2AX alongside IdU that peaks before declining below its baseline value, while PDO27 has a decrease in pRB.

**Fig. B2.**
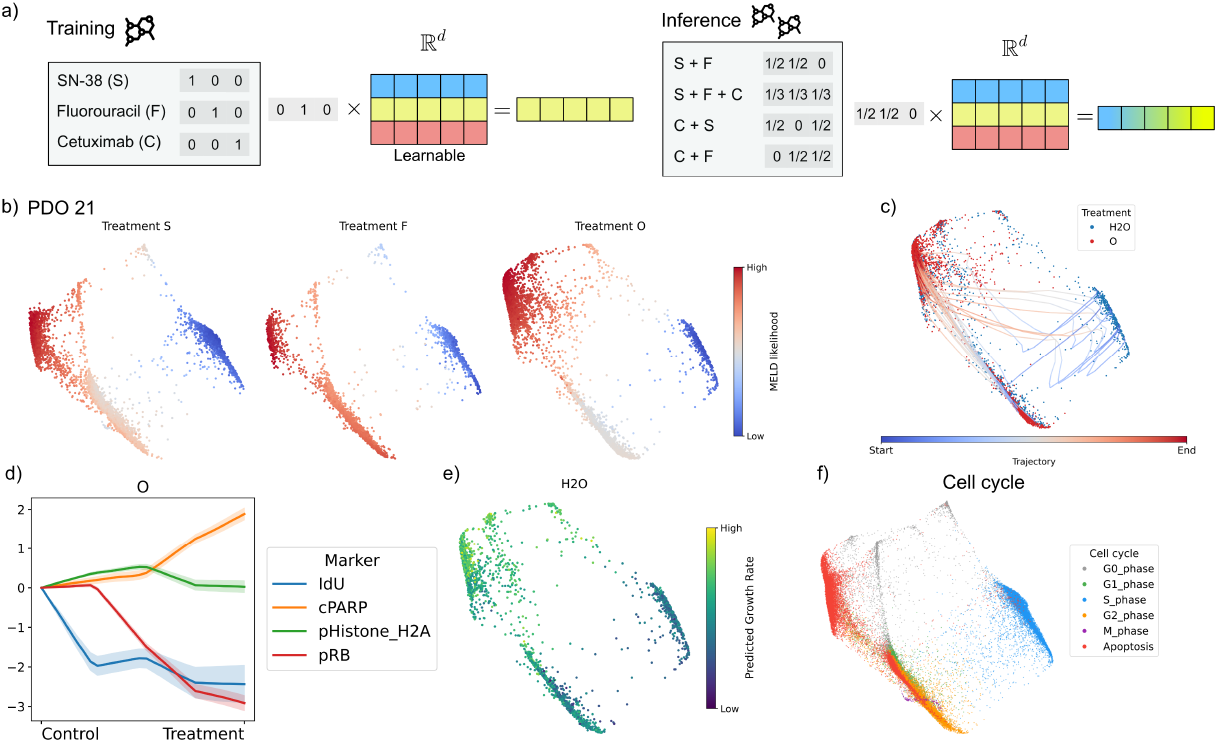
GeoSinkhorn Flow trajectories and growth-rate inference for PDO21. a) Training and inference pipelines for compositional treatment encoding of single and combinatorial perturbations. b) MELD likelihood values across cells for single treatments (S, F, and O). c) Inferred cellular trajectories in PHATE space for treatments O. d) Predicted marker-level trajectories for treatment O evaluated on apoptosis and cell-cycle markers (pRB, cPARP, pHistone H2A, IdU). Values show mean marker changes relative to the control condition at *t* = 0. e) Predicted growth rate across H_2_O control cells over PHATE. f) Cell-cycle phase over PHATE.

The consistency of trajectory start and end points was assessed for PDO 21 by computing Pearson and Spearman correlations between predicted and ground-truth trajectory start and end points, for both seen single-drug and unseen combinatorial treatment conditions (Supplementary Figure B5). For seen single-drug treatments, the model demonstrated consistently high agreement at trajectory start points, with mean Pearson and Spearman correlations of 0.996 and 0.988 respectively, indicating near-perfect linear and rank-order correspondence. End point correlations remained strong, though slightly reduced (mean Pearson: 0.894, Spearman: 0.854), which is expected given the greater uncertainty in predicting terminal states. Generalization to unseen combinatorial treatments followed a similar pattern, with start point correlations of 0.894 (Pearson) and 0.952 (Spearman), and end point correlations of 0.835 (Pearson) and 0.876 (Spearman). Notably, the one outlying value of 0.499 visible in the unseen condition represents an isolated case and does not reflect broader model performance.

**Fig. B3.**
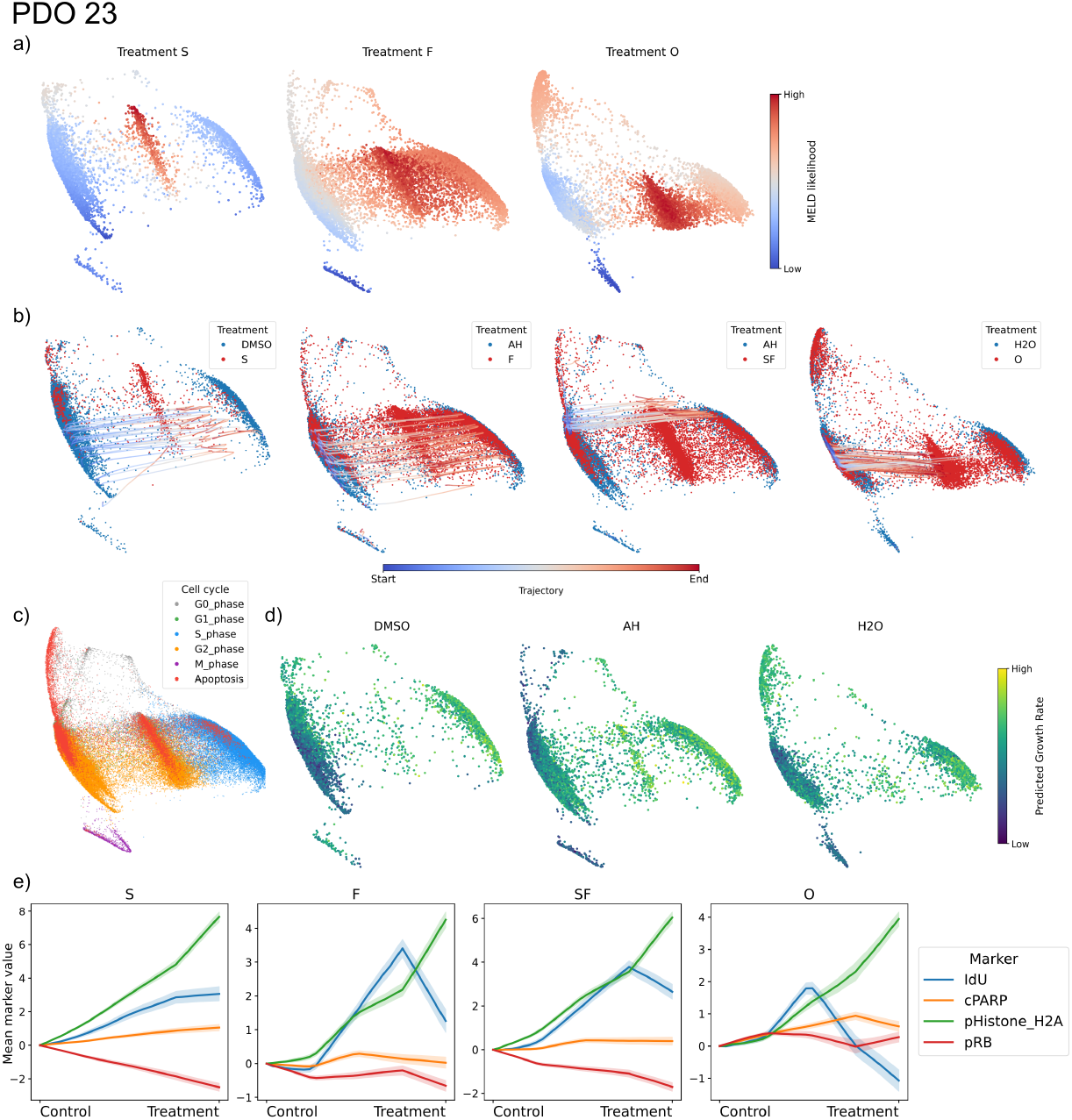
GeoSinkhorn Flow trajectories and growth-rate inference for PDO23. a) MELD likelihood values across cells for single treatments (S, F, and O). b) Inferred cellular trajectories in PHATE space for single treatments (S, F, O) and the held-out combination treatment SF. c) Cell-cycle phase over PHATE. d) Predicted growth rate across control cells over PHATE. e) Predicted marker-level trajectories for single treatments (S, F, O) and held-out combinations (SF) evaluated on apoptosis and cell-cycle markers (pRB, cPARP, pHistone H2A, IdU). Values show mean marker changes relative to the control condition at *t* = 0.

The consistently higher Spearman relative to Pearson correlations in the unseen setting suggests that while absolute value predictions introduce some noise, the model preserves the correct ordering of perturbation responses even under distributional shift.

#### B.4 Perturb-seq additional details and results

##### Perturb-seq cell and gene filtering

The Perturb-seq expression matrix was processed from the aggregated gene-by-cell count matrix and guide-assignment table. The preprocessed AnnData object contained 214,449 cells measured over 36,601 genes. Mitochondrial, ribosomal and haemoglobin genes were annotated for quality-control summaries, and per-cell QC metrics were computed using Scanpy. The expression matrix was library-size normalized with `scanpy.pp.normalize total` and log-transformed with `scanpy.pp.log1p`.

**Fig. B4.**
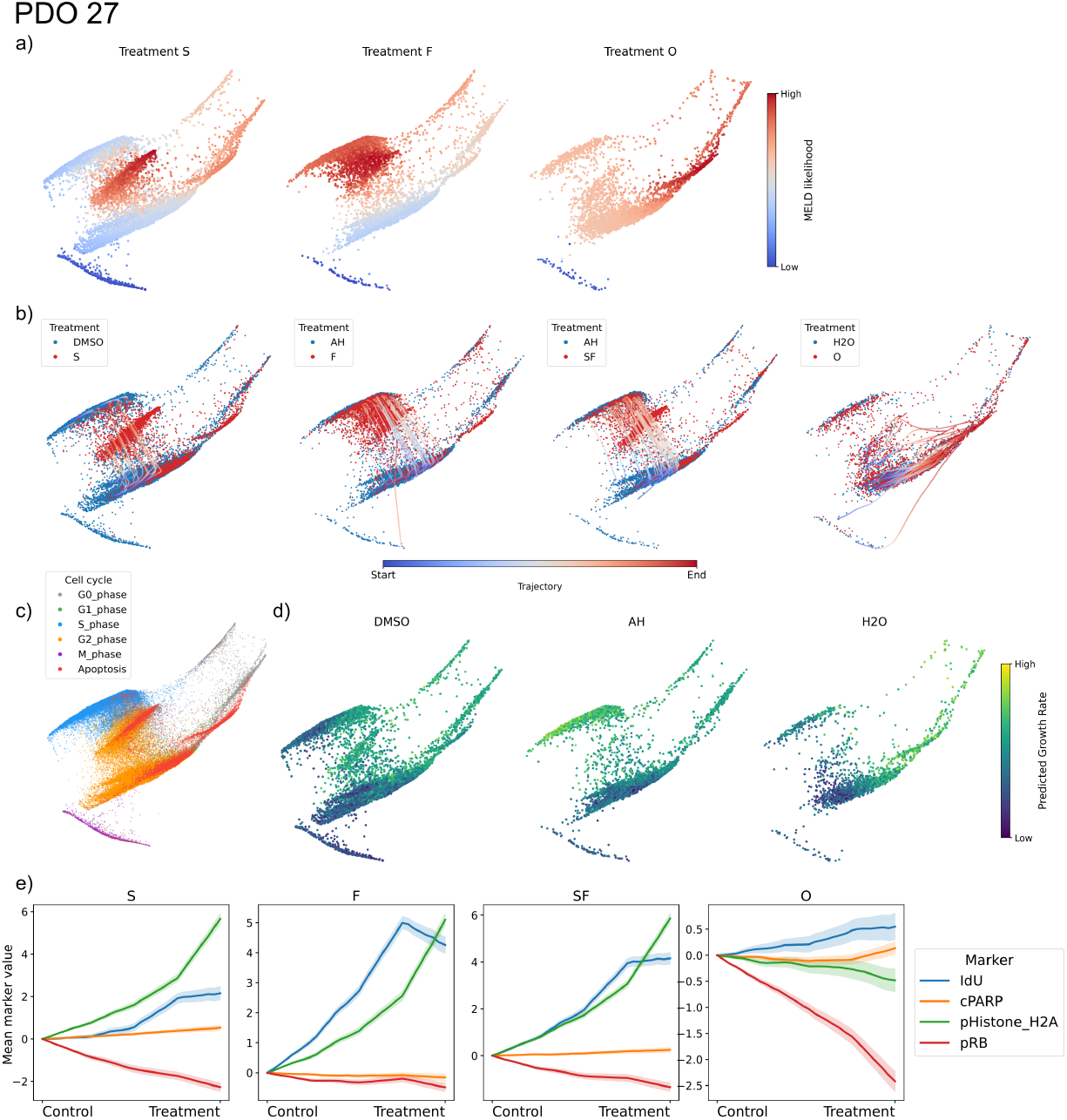
GeoSinkhorn Flow trajectories and growth-rate inference for PDO 27. a) MELD likelihood values across cells for single treatments (S, F, and O). b) Inferred cellular trajectories in PHATE space for single treatments (S, F, O), and combination treatment (SF). c) cell-cycle phase over PHATE. d) Predicted growth rate across control cells over PHATE. e) Predicted marker-level trajectories for single treatments (S, F, O) and held-out combination (SF) evaluated on apoptosis and cell-cycle markers (pRB, cPARP, pHistone H2A, IdU). Values show mean marker changes relative to the control condition at *t* = 0.

For the Perturb-seq experiments, we selected the top 5,000 highly variable genes using Scanpy with `n top genes=5000`. Batch effects were corrected on this 5,000-gene matrix using ComBat, with experimental batch as the covariate. The resulting batch-corrected AnnData object contained 214,449 cells and 5,000 genes, and was used as the input expression space for the Perturb-seq autoencoder and downstream flow model.

For the experiment, perturbation labels were filtered at the condition level. We retained perturbations with more than 32 cells, required an available GeneWalk embedding, and restricted the analysis to a curated CAD/endothelial perturbation set. This selected set contained 33 non-control perturbations together with the *negative-control* group. Six CAD-relevant perturbations were withheld from training to evaluate generalization to unseen perturbation genes: *CCM2, MAP2K5, EDN1, NOS3, TSPAN14* and *PECAM1*. The remaining 27 non-control perturbations were used for training.

**Fig. B5.**
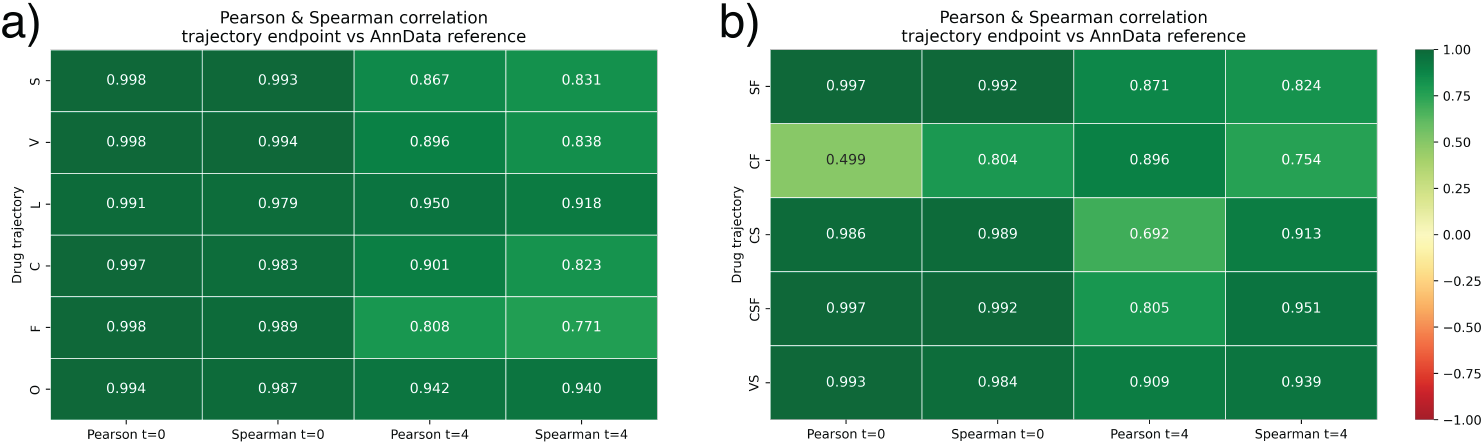
Correlation analysis of trajectory start and end points in the PDO dataset. (a) Pearson and Spearman correlations for seen single-perturbation data; (b) for unseen combinatorial perturbation data.

**Fig. B6.**
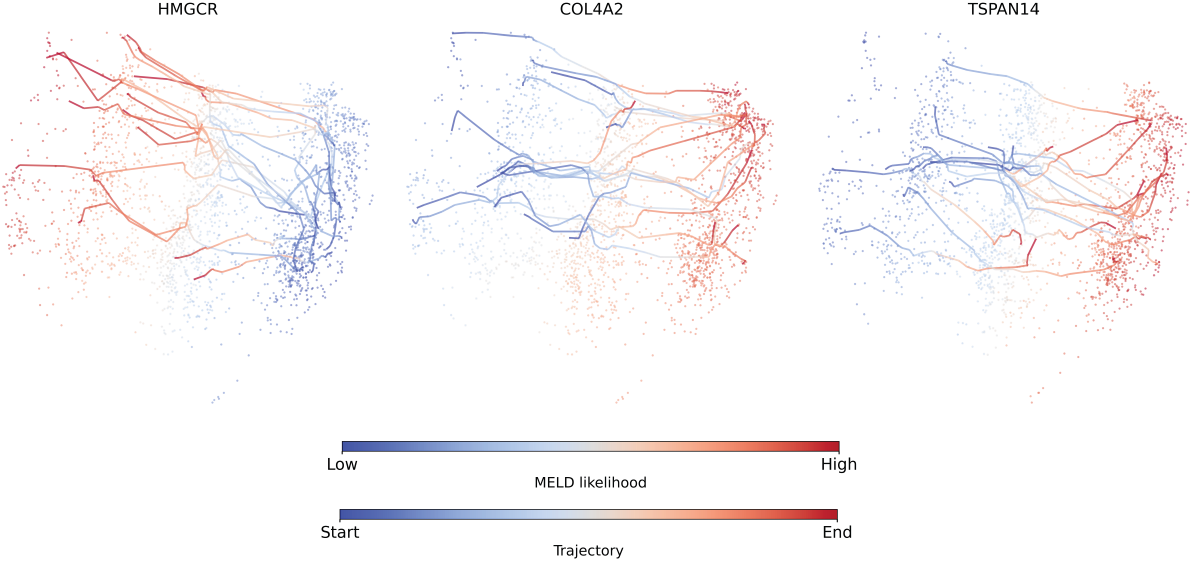
Pair-specific MELD likelihoods and inferred GeoSinkhorn Flow trajectories for representative CAD-related perturbations. Each panel shows the cell-level PHATE embedding for one control-to-perturbation comparison. Cells are colored by MELD likelihood from control-like to perturbation-like states, and overlaid paths show inferred trajectories colored from start to endpoint. *HMGCR* and *COL4A2* were included in training, whereas *TSPAN14* was held out for unseenperturbation evaluation.

The final training object contained 4,660 cells, comprising 2,438 perturbed cells and 2,222 negative-control cells. The held-out test object contained 2,754 cells, comprising 532 held-out perturbed cells and the same 2,222 negative-control cells used as the common source population. The training perturbations were *CCL2, CDH5, CDKN2B, COL4A1, COL4A2, CTNNA1, EGFL7, ENG, FLI1, FLT1, HEG1, HMGCR, HTRA1, JAK1, JUN, KLF4, KRIT1, LOX, MMP1, NRP1, NRP2, PDGFB, POSTN, SMAD3, SNAI2, SOX4* and *THBS1*.

#### B.5 CyTOF signaling additional details and results

The main CyTOF signaling figure highlights three representative PI3K/MAPK read-outs. To show the remaining model outputs, Figure B7a reports generated trajectories for the five markers not displayed in the main panel: pNDRG1^T346^, pSMAD2/3, total S6, pRB^S807*/*S811^ and cleaved caspase-3^D175^. These trajectories provide an additional marker-level view of the generated signaling dynamics across *PIK3CA* genotype, stimulation condition and time.

As a distribution-level check, Figure B7b compares generated and original CyTOF marker distributions for each genotype–treatment–time condition. For each marker and condition, the one-dimensional Wasserstein distance was computed between the generated and original single-cell marker distributions after the same preprocessing used for model training. Smaller values indicate closer agreement between generated and original distributions.

**Fig. B7.**
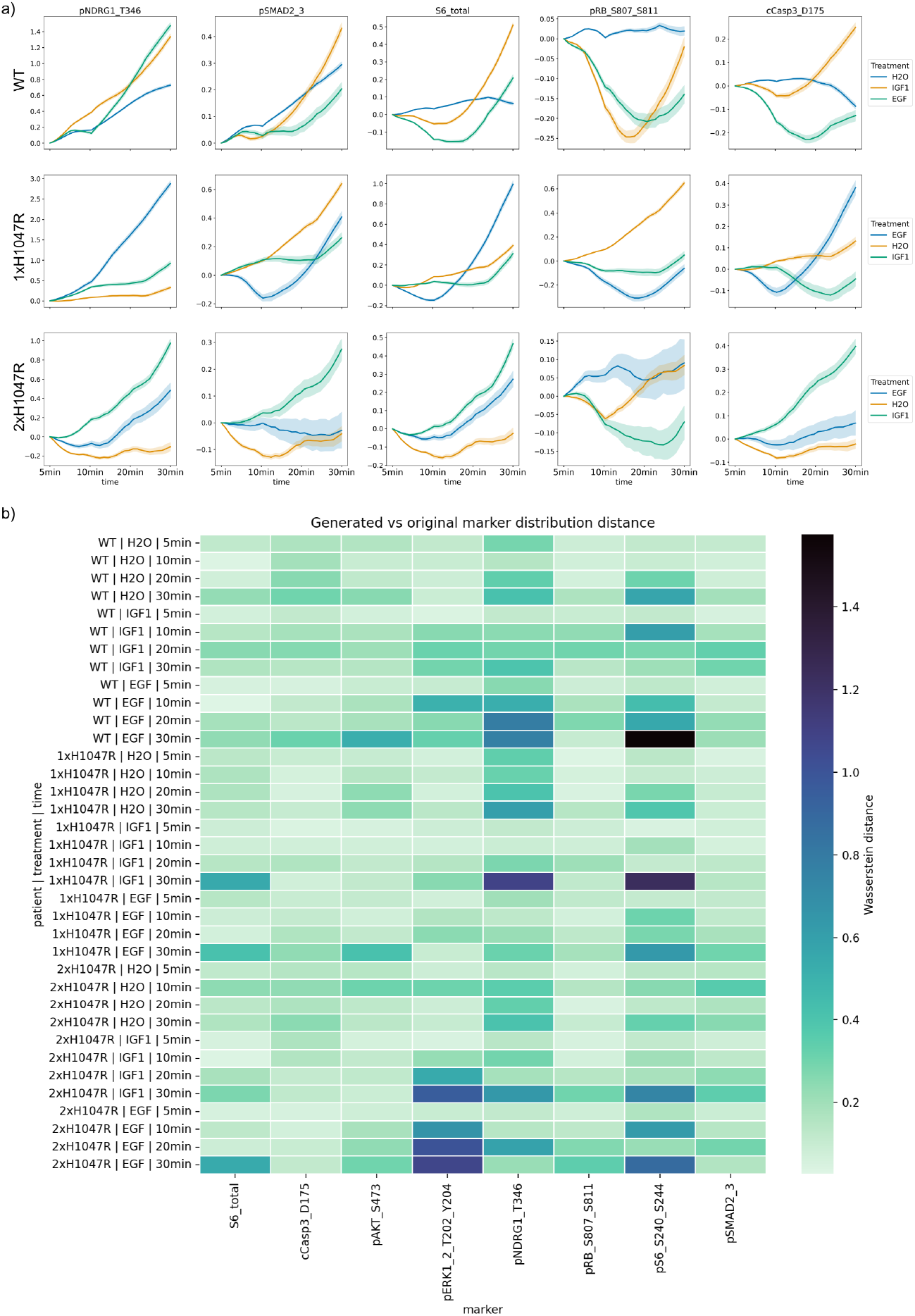
a) Generated marker-level trajectories for the five CyTOF markers (Section 4.8.4) not shown in the main figure. Trajectories are shown across *PIK3CA* genotype, stimulation condition and time. b) Marker-wise Wasserstein distances between generated and original single-cell marker distributions for each genotype–treatment–time condition. Lower values indicate closer agreement between generated and original distributions.

## References

[1] Qin, X. et al. An oncogenic phenoscape of colonic stem cell polarization. Cell 186, 5554–5568.e18 (2023).

[2] Huguet, G. et al. Geodesic sinkhorn for fast and accurate optimal transport on manifolds, 1–6 (IEEE, 2023).

[3] Huguet, G. et al. A heat diffusion perspective on geodesic preserving dimensionality reduction. arXiv preprint arXiv:2305.19043 (2023).

[4] Zapatero, M. R. et al. Trellis single-cell screening reveals stromal regulation of patient-derived organoid drug responses. bioRxiv (2023).

[5] Burkhardt, D. B. et al. Quantifying the effect of experimental perturbations at single-cell resolution. Nature biotechnology 39, 619–629 (2021).

[6] Schnitzler, G. R. et al. Convergence of coronary artery disease genes onto endothelial cell programs. Nature 626, 799–807 (2024).

[7] Madsen, R. R. et al. Oncogenic pik3ca corrupts growth factor signaling specificity. Molecular Systems Biology 21, 126–157 (2025).

[8] Moon, K. R. et al. Visualizing structure and transitions in high-dimensional biological data. Nat Biotechnol 37, 1482–1492 (2019).

[9] Solomon, J. et al. Convolutional wasserstein distances: Efficient optimal transportation on geometric domains. ACM Transactions on Graphics (ToG) (2015).

[10] Varadhan, S. R. S. On the behavior of the fundamental solution of the heat equation with variable coefficients. Communications on Pure and Applied Mathematics (1967).

[11] Shuman, D. I., Vandergheynst, P. & Frossard, P. Chebyshev polynomial approximation for distributed signal processing (IEEE, 2011).

[12] Marcotte, S. et al. Fast multiscale diffusion on graphs (IEEE, 2022).

[13] Varadhan, S. R. S. On the behavior of the fundamental solution of the heat equation with variable coefficients. Communications on Pure and Applied Mathematics 20, 431–455 (1967).

[14] Lipman, Y., Chen, R. T. Q., Ben-Hamu, H., Nickel, M. & Le, M. Flow matching for generative modeling (2023). arXiv:2210.02747.

[15] Tong, A. et al. Improving and generalizing flow-based generative models with minibatch optimal transport (2023). arXiv:2302.00482.

[16] Ietswaart, R., Gyori, B. M., Bachman, J. A., Sorger, P. K. & Churchman, L. S. Genewalk identifies relevant gene functions for a biological context using network representation learning. Genome biology 22, 55 (2021).

[17] Cuturi, M. Sinkhorn distances: Lightspeed computation of optimal transport. NeurIPS (2013).

[18] Peyré, G. & Cuturi, M. Computational optimal transport: With applications to data science (Now Foundations and Trends, 2019).

[19] Benamou, J.-D. & Brenier, Y. A computational fluid mechanics solution to the Monge-Kantorovich mass transfer problem. Numerische Mathematik 84, 375–393 (2000).

[20] Paszke, A. et al. Pytorch: An imperative style, high-performance deep learning library. Advances in neural information processing systems 32 (2019).

[21] Falcon, W. & team, T. P. L. Pytorch lightning. Zenodo (2019). URL https://github.com/Lightning-AI/pytorch-lightning.

[22] Yadan, O. Hydra: A framework for elegantly configuring complex applications. GitHub (2019). URL https://github.com/facebookresearch/hydra.

[23] Flamary, R. et al. POT: Python optimal transport. Journal of Machine Learning Research 22, 1–8 (2021). URL https://jmlr.org/papers/v22/20-451.html.

[24] Tong, A. et al. Improving and generalizing flow-based generative models with minibatch optimal transport. Transactions on Machine Learning Research (2024). URL https://openreview.net/forum?id=CD9Snc73AW.

[25] Tong, A. Y. et al. Meila, M. & Zhang T. (eds) Diffusion earth mover’s distance and distribution embeddings. (eds Meila, M. & Zhang,T.) Proceedings of the 38th International Conference on Machine Learning, Vol. 139 of Proceedings of Machine Learning Research, 10336–10346 (PMLR, 2021). URL http://proceedings.mlr.press/v139/tong21a.html.

